# Transcriptomic analyses reveal prenatal stress promotes late-onset neural stem cell proliferation in adult male offspring

**DOI:** 10.1101/2022.12.15.520572

**Authors:** Z. Wang, L.Q. Zhou, C.M. Lee, J.Y. Shi, L. Liu, X.J. Yang, Y.X. Deng, J.P. Liu, J.B. Wang, W.M. Zhu, Y.E. Sun, Q. Lin

**Affiliations:** Shanghai Institute of Stem Cell Research and Clinical Translation, Shanghai East Hospital, School of Medicine, Tongji University, Shanghai, China; Department of Psychiatry and Behavioral Sciences, Intellectual Development and Disabilities Research Center, University of California, Los Angeles, CA 90095; Stem Cell Translational Research Center, Tongji Hospital, School of Medicine, Tongji University, Shanghai, China

**Author notes:** To whom correspondence may be addressed. Q.L., Y.E.S., or. Equal contribution (Z. Wang, L.Q. Zhou, C.M. Lee). Des Moines University, 3200 Grand Ave. Des Moines, IA 50312.

## Abstract

Perturbations during critical time windows during embryonic development can lead to adverse functional consequences that manifest later in life. Here, we report that prenatal maternal stress (PNS) during the peak of embryonic neurogenesis (E14-delivery) dramatically increased numbers of proliferating neural stem/progenitor cells (NSC/NPCs) in the ependymal-subventricular zone (E-SVZ) and neuroblasts (NBs) in the rostral migratory stream and newborn neurons in the olfactory bulb (OB) of male mouse offspring without causing significant cell death or a deficit in cell migration to the OB. Mechanistically, bulk and single-nucleus transcriptomic analyses showed that PNS affected gene regulatory networks controlling cell cycle progression and stem cell maintenance, maturation of neural circuits, and gliogenesis in prenatally stressed (STR) offspring. More specifically, we found that prenatal exposure to mild maternal restraint-stress sustained MAPK (ERK) activity in the E-SVZ and thus prolonged NSC/NPC/NB proliferation in stressed brains. Moreover, we found PNS disorganized the cytoarchitecture of the glomerular layer of the OB, which may directly relate to the deficit in discriminating different social orders in stressed offspring. Compared to STR males, their female littermates showed less change in the number of proliferating cells in the E-SVZ.

## Introduction

In humans and model animals, environmental disturbances during the critical period have been shown to cause neural morphological, neurobiological, and behavioral deficits [1-11]. Since the evidence of cell mitosis in the SVZ of adult rats was published over 100 years ago [12], it has now been widely accepted that adult neurogenesis occurs in mammals [13-17]. Numerous studies have reported that the ependymal-subventricular zone (E-SVZ) and the dentate gyrus (DG) are the brain regions that harbor NSC/NPCs to give rise to neurons and glial cells throughout life [14, 15, 18-22]. Mechanistically, epidermal growth factor (EGF) and vascular endothelial growth factor (VEGF) are two key extrinsic factors to activate dormant/quiescent neural stem cell and neural progenitor cell proliferation in the E-SVZ of the ventricles in mouse brain [14, 19]. MAPK (a.k.a. extracellular signal-regulated kinase (ERK)) cascade, as a major transducer of growth factors like EGF and VEGF, plays a vital role in mediating their intracellular signaling [23, 24]. MAPK(ERK) has a TxY motif in its activation domain, which requires dual phosphorylation by upstream MAPK kinases (e.g., MAP2K1/2 (MEK1/2)) for catalytic activation. MAPK3 and 1 (ERK1/2) have been shown to interact with wide-range signaling pathways, such as EGF/FGF/VEGF, WNT, BMP/TGFB, Estrogen, and SHH [25-29] to regulate cell proliferation and to facilitate epithelial-mesenchymal transition (EMT) in cancer cells [30, 31].

Although tremendous progress has been made to understand the long-lasting effects of PNS on brain development, little is known about the molecular programs that govern adult NSC/NPC proliferation and neurogenesis in response to prenatal stimuli. In the current study, we found that mild, prenatal maternal restraint stress increased the numbers of BrdU^+^ and MKI67^+^ proliferating cells in the E-SVZ and the rostral migratory stream (RMS) and the number of BrdU^+^/RBFOX3^+^ postmitotic neurons in the OB. This finding contradicted the normal finding that prenatal adverse stimulus impaired neurogenesis in postnatal brains [32-35].. To explore the basic molecular program that PNS employed to maintain stem cell proliferation in adult brains, we carried out bulk and single-nucleus RNAseq using E-SVZ tissues collected from control and stressed mouse brains. Gene regulatory network analysis indicated that PNS dysregulated the expression of genes that regulate cell cycle progression and neural stem cell maintenance, synapse organization and membrane potential, and gliogenesis. Particularly, we showed that the mild maternal stress at the mid-late stage of gestation sustained postnatal phosphorylation levels of MAPK3/1 in the E-SVZ, which in turn promoted NSC/NPC proliferation in adult male brains. This finding suggests that MAPK (ERK) is a bona fade downstream signaling cascade mediating extrinsic cues to stimulate cell proliferation in the adult brain. Furthermore, we found that prenatally stressed male mice had an intact olfactory function to locate food pellets and distinguish nonsocial odors, whereas they showed impairment in discriminating different social smells.

## Results

### PNS triggers cell proliferation in the lateral E-SVZ

To evaluate the effects of the prenatal aversive stimulus on cell proliferation in the E-SVZ, we employed a widely used stress-inducing method, the restraint stress on pregnant dams during the third week of gestation (60min/day, 0800h-0900h) [36]. The third week of gestation in mice roughly corresponds to the second trimester of human gestation, which is a critical period for the development of psychiatric disorders [37, 38]. To eliminate the postnatal influence from stressed mothers, the prenatally stressed (STR) pups were fostered by unstressed dams at postnatal day 0 (P0). As a control (CTL), pups from unstressed dams also underwent the fostering process. BrdU was administered by intraperitoneal (IP) injection one hour before tissue collection (Fig.1A).

**Fig. 1.**
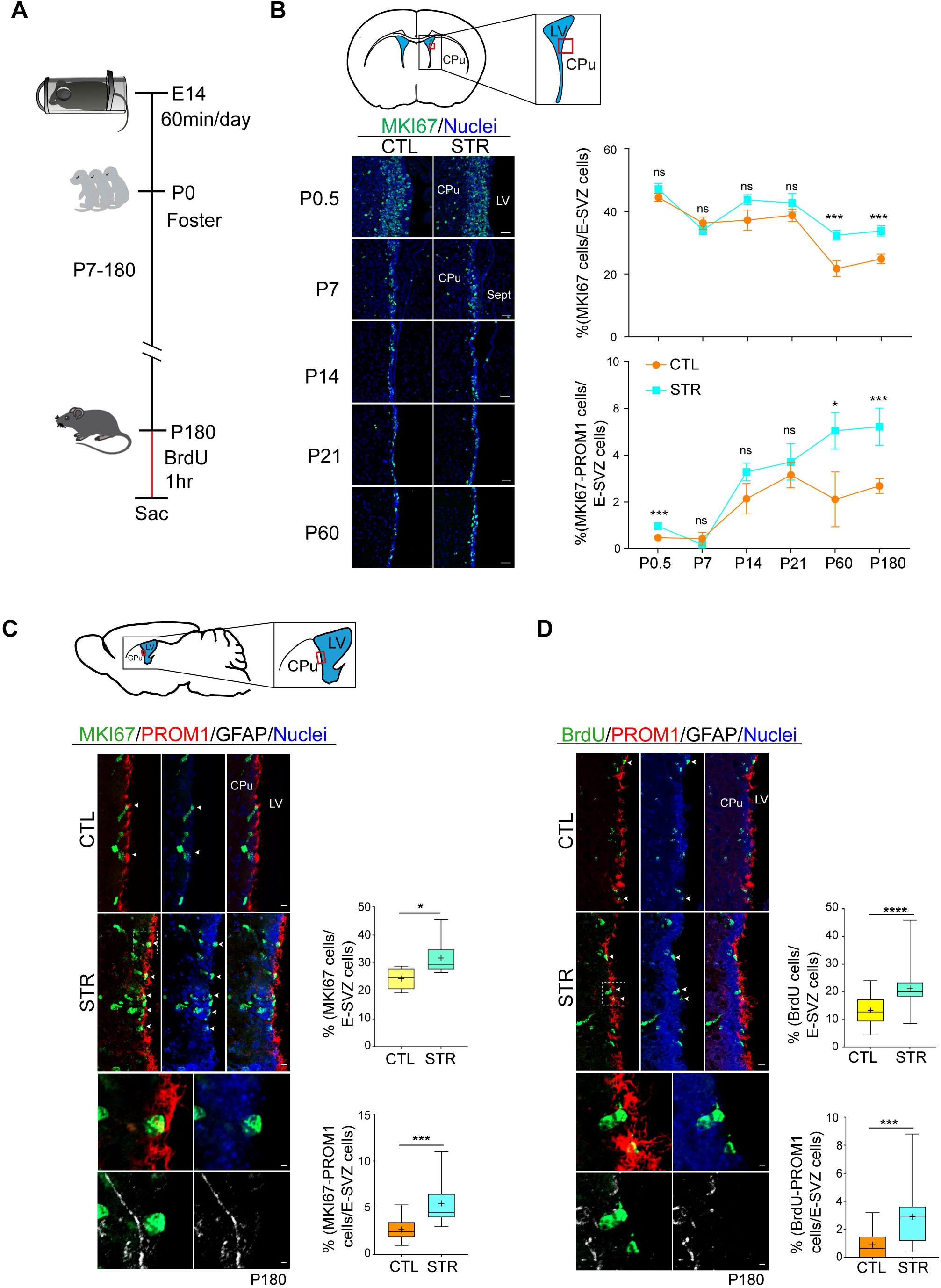
PNS promoted PROM1^+^ cell proliferation in the lateral E-SVZ of prenatally stressed male offspring. **(A)** The pregnant dams were stressed in a plastic restrainer from E14 until delivery. Samples were collected at postnatal different time points. To label proliferating cells, BrdU was IP injected 1hr before sacrifice. **(B)** PNS significantly increased E-SVZ NSC proliferation in prenatally stressed adult offspring (P60 and P180), but not neonatal and juvenile mice (P0.5-21). CPu, caudate putamen; LV, lateral ventricle; Sept, septum. Scale bar, 30 μm. **(C)** MKI67 (green color), PROM1 (red color), and GFAP (white color) immunostaining. The enlarged images at the bottom showed MKI67^+^/PROMl^+^ ependymal cells are GFAP negative (MKI67^+^ cells: CTL = 24.5 ± 1.6%, n = 6; STR = 31.8 ± 2.8%, n=6. P = 0.026. MKI67^+^/PROM1^+^ cells: CTL = 2.7 ± 0.3%, n = 13; STR= 5.5 ± 0.8%, n = 10. P = 0.0002). The box plot showed the minimum, the maximum, the median, the mean (the + symbol), and the first and the third quartiles. Scale bars, 10 μm, 2 μm. **(D)** BrdU (green color), PROM1 (red color), and GFAP (white color) immunostaining (BrdU^+^ cells: CTL = 13.3 ± 0.9%, n = 28; STR =21.4 ± 1.4%, n = 26. P<0.0001. BrdU^+^/PROM1^+^ cells: CTL = 0.9 ± 0.2%, n = 26; STR = 2.9 ± 0.5%, n = 18. P = 0.0002). The enlarged images on the bottom showed BrdU/PROMl double positive ependymal cells were GFAP negative. Scale bars, 10 μm, 2 μm. CTL, control; STR, stressed. Mann-Whitney test. All data were presented as mean ± SEM. *p < 0.05, ***p < 0.001, ****p < 0.0001. For sample quantification, we examined sections collected from minimal 3-5 brains of different litters per experimental condition.

Compared to the CTL, in STR male offspring the ratio of proliferating cells to the total number of cells in the lateral E-SVZ (MKI67^+^ cells/total E-SVZ cells) was dramatically increased at P60 and P180, whereas similar increases were not detected in neonatal or juvenile mice (P0.5-21) (Fig.1B; Sup.Fig.1). We found that the numbers of BrdU^+^/PROM1^+^/GFAP^-^ and MKI67^+^/PROM1^+^/GFAP^-^ ependymal cells, and MKI67^+^/DCX^+^, BrdU^+^/DCX^+^, and MKI67^+^/TUBB3^+^ neuroblasts increased dramatically in STR male mice (Figs.1C, D, 2A; Sup.Fig.3A). This finding is in line with a previous study that showed ‘mild’ prenatal stress (30min/day, E15-17) enhanced neurogenesis in the DG of neonatal rat brains [39]. There were more BrdU^+^ and MKI67^+^ migrating neuroblasts in the rostral migratory stream (RMS) of stressed male mice than that of the CTL mice (Fig.2B). We did not observe statistically significant changes in the numbers of GFAP^+^/BrdU^+^ cells in either female or male offspring (Sup.Figs.2D, 3B).

**Fig. 2.**
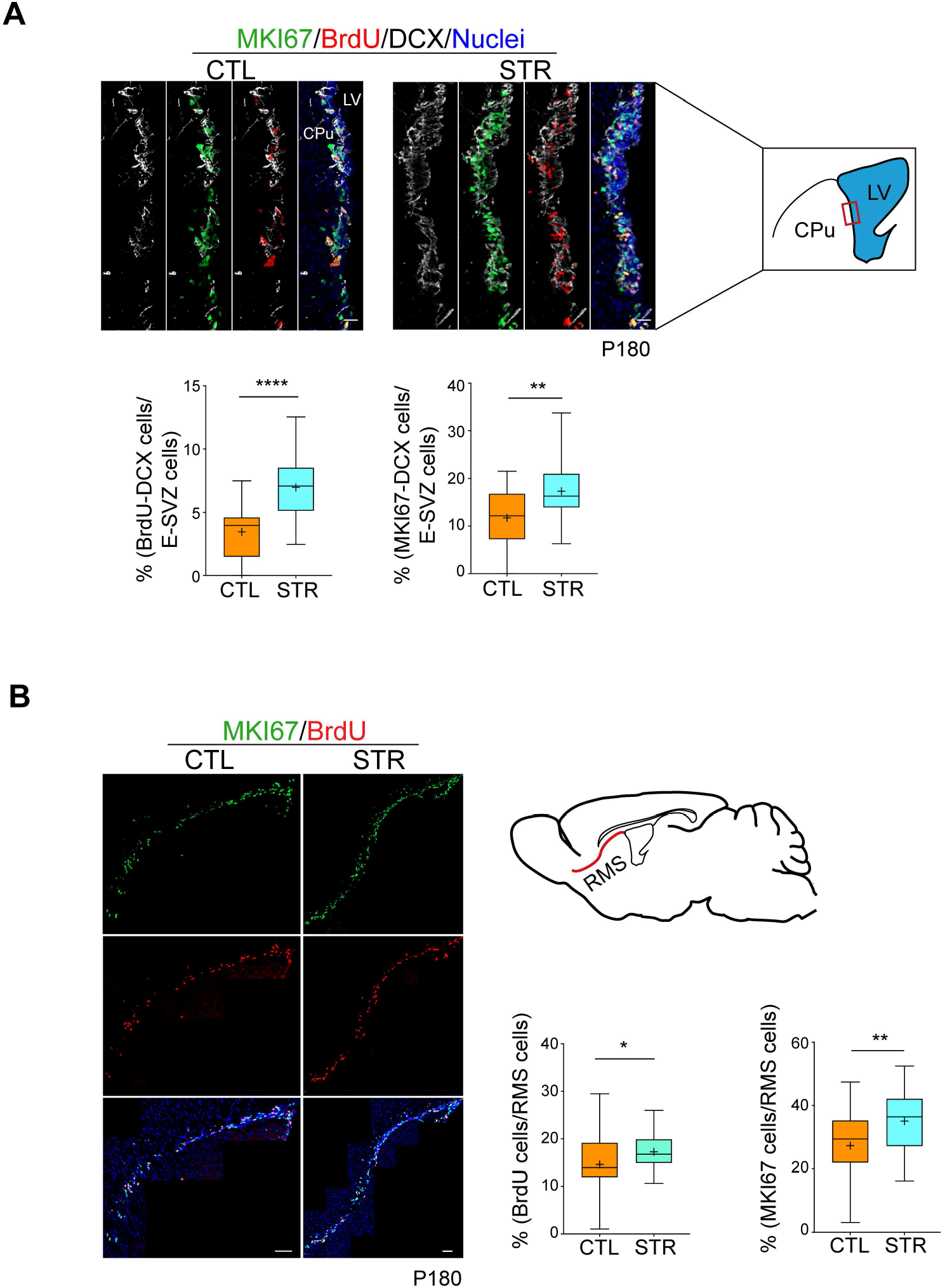
PNS promotes neuroblast proliferation in the E-SVZ and the RMS of male offspring. **(A)** MKI67/BrdU/DCX immunostaining shows that the ratio of neuroblasts increased dramatically in the lateral E-SVZ (BrdU^+^/DCX^+^ cells: CTL = 3.5 ± 0.4%, n = 21; STR = 7.0 ± 0.6%, n = 23. P<0.0001. MKI67^+^/DCX^+^ cells: CTL = 11.8 ± 1.3%, n = 21; STR = 17.3 ± 1.3%, n = 23. P = 0.0071). Scale bars, 30 μm. **(B)** The ratio of proliferating A-cell (BrdU^+^ or MKI67^+^) increased in the RMS (MKI67^+^ cells: CTL = 27.3 ± 1.8%, n = 38; STR = 35.1 ± 1.4%, n = 40. P = 0.0021. BrdU^+^ cells: CTL = 14.6 ± 1.1%, n = 38; STR = 17.3 ± 0.6%, n = 40. P = 0.021). Scale bars, 100 μm. Mann-Whitney test. All data were presented as mean ± SEM. *p < 0.05, **p < 0.01, ****p < 0.0001.

In the female littermates, the changes were statistically insignificant, suggesting that female offspring were resilient to PNS (Sup.Figs.2, 3C,D). We found that under normal physiological conditions, the baseline of the ratio of proliferating cells to total cells in the E-SVZ in female mice was higher than that of their male littermates (BrdU^+^/PROM1^+^ cells: female = 2.1 ± 0.4% vs. male = 0.9 ± 0.2%; MKI67^+^/PROM1^+^ cells: female = 4.1 ± 0.4% vs. male = 2.7 ± 0.3%. mean ± SEM). A similar finding was demonstrated in the subgranular zone of the hippocampal dentate gyrus in female mice by Tanapat et al [40]. Moreover, studies have shown that the influence of PNS exposure on the offspring is highly sex dependent. In rats, PNS significantly increased anxiety- and depression-related behaviors in male offspring, but not in female offspring [41, 42]. In humans, it has been shown that prenatal disturbances preferentially affect male offspring in a late-onset manner [43, 44].

### E-SVZ NSC/NPC progenies migrate to and mature in the olfactory bulb (OB)

To observe bulbar neurogenesis in the STR offspring, we labeled proliferating cells with two BrdU administrations separated by a 12-hr interval. The animals were sacrificed six weeks later (Fig.3A). BrdU-labeled cells migrated to the OB and mainly resided in the granule cell layer (GrO) with a few cells in the glomerular layer (Gl) and the external plexiform layer (EPl) in both CTL and STR offspring (8-9-month-old). As expected, the number of BrdU-labeled, newborn RBFOX3^+^ neurons increased in the OB of stressed mice (Fig.3B, C). We did not observe substantial cell death in the OB of either the CTL or STR mice (Fig.3D).

**Fig. 3.**
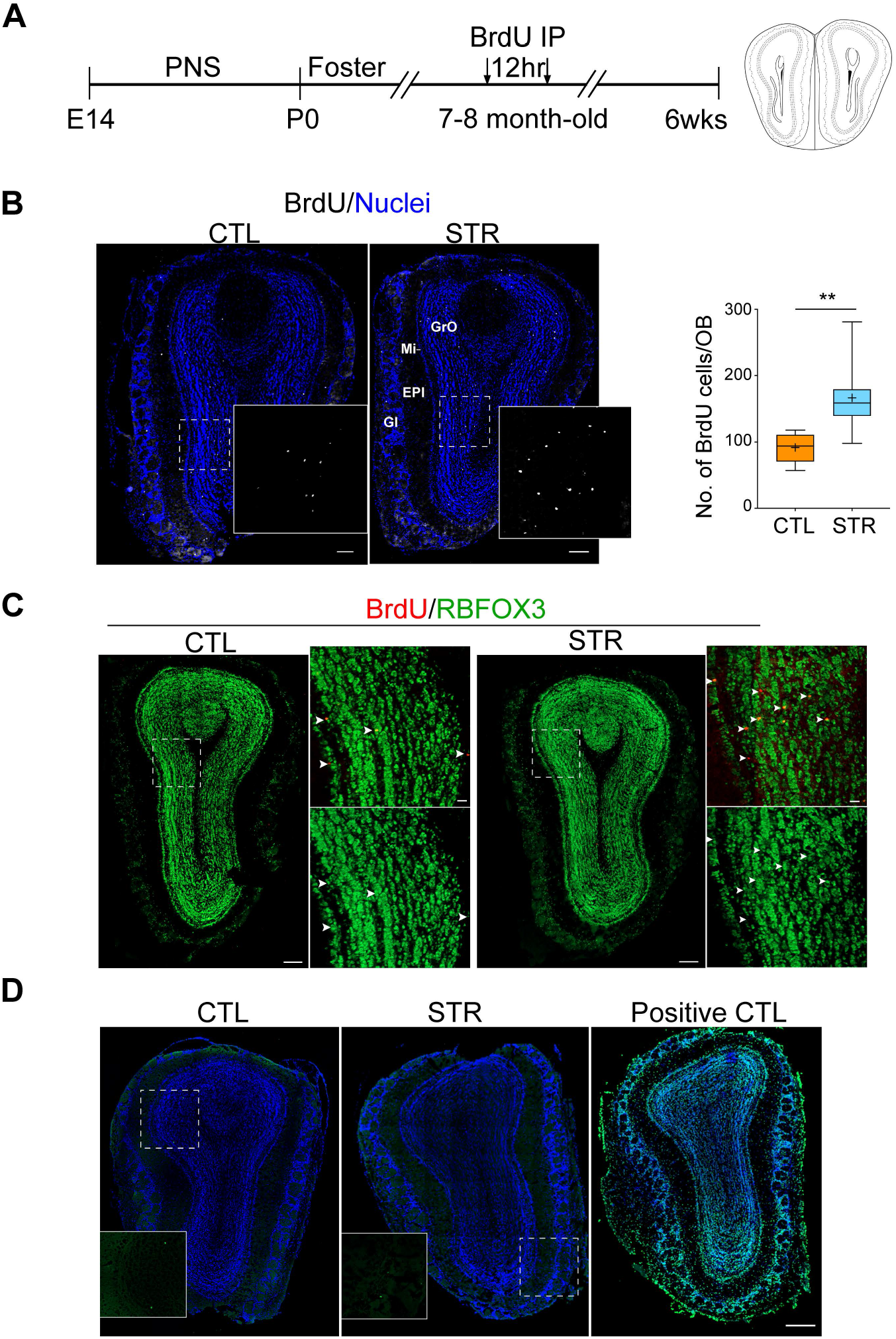
PNS disturbs bulbar neurogenesis. **(A)** To determine fates of lateral E-SVZ NSCs in prenatally stressed male mice, NSCs were labeled with two BrdU IP injections. After 6 weeks, the brain samples were collected. **(B)** BrdU immunostaining (white color). The enlarged images on the right show BrdU staining in the GrO (BrdU^+^ cells/OB section: CTL = 92 ± 8, n = 8. STR = 166 ± 19, n =8. P = 0.0011. **p < 0.01. mean ± SEM). Mann-Whitney test. Scale bar, 200 μm. **(C)** BrdU labeled cells migrated to the OB and matured to become RBFOX3^+^ neurons. The arrowheads show BrdU/RBFOX3 double-positive cells in the GrO. Scale bars, 200 μm. **(D)** TUNEL assay shows that PNS does not cause cell apoptosis in the OB. Scale bars, 300 μm. GrO, granular cell layer of OB; Mi, mitral cell layer; EPl, external plexiform layer; Gl, glomerular layer.

### Bulk and single nucleus RNAseq (snRNAseq) analyses show that PNS promotes E-SVZ cell proliferation via upregulating MAPK (ERK) pathway

In this study, we showed mild prenatal restraint stress enhanced the proliferation potency of NSC/NPCs in adult STR offspring. This finding contradicted the normal finding in the field that prenatal adverse stimulus impaired neurogenesis in postnatal brains [32, 33], suggesting that different stress levels ranging from mild to severe can have different negative influences on the neural development of offspring. To reveal the long-lasting molecular regulation that PNS elicited an increase in cell proliferation, we carried out transcriptomic analyses. Bilateral E-SVZ regions from 49 eight-month-old STR male and female mice were collected, rapidly frozen using liquid nitrogen, and stored at -80°C. Tissues from the left and right sides were stored separately: one for bulk RNAseq, and one for snRNAseq (Fig.4A).

**Fig. 4.**
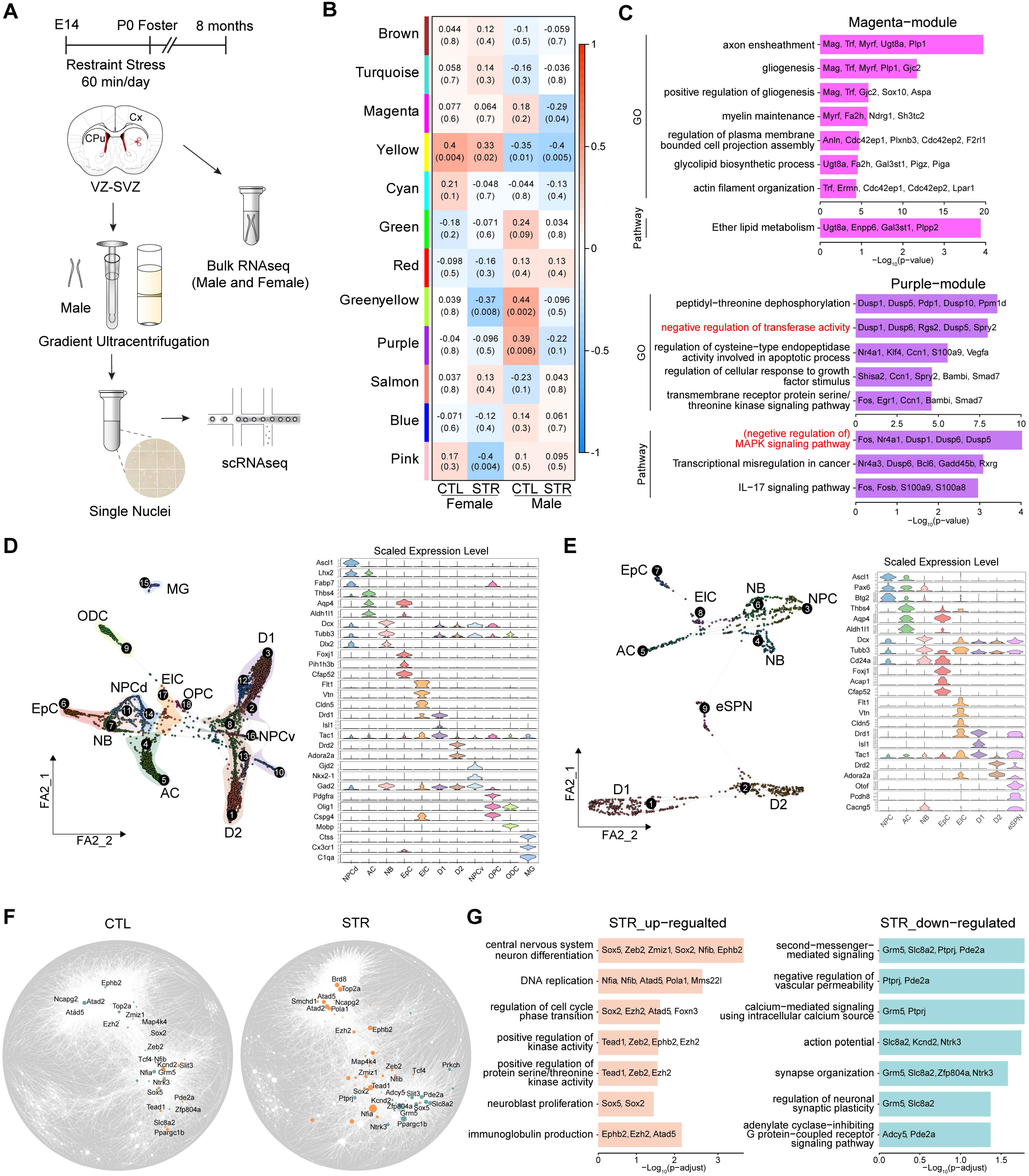
Bulk and single nucleus transcriptomic analyses of cells in the lateral E-SVZ of STR and CTL mice. **(A)** Schematic overview of tissue and single nucleus isolation for bulk RNA sequencing and single nucleus RNA sequencing (snRNAseq). **(B)** Module-trait correlation analysis revealed 3 modules in male offspring were correlated with PNS (Magenta, Greenyellow, Purple). **(C)** GO analysis shows top 7 GO terms enriched in major gene modules (Magenta and Purple), excluding redundant terms. The top 5 enriched genes of each term are listed. **(D)** Force-directed graph is used to visualize clusters of the total single nucleus dataset (the left panel). EpC, ependymal cell; ElC, endothelial-like cell; NPCd, dorsal neural progenitor cell; NPCv, ventral neural progenitor cell; NB, neuroblast; D1, dopamine receptor D1, Drd1^+^ spiny projection neuron; D2, Drd2^+^ spiny projection neuron; AC, astrocyte; OPC, oligodendrocyte progenitor cell; OD, oligodendrocyte; MG, microglia. The violin plot (the right panel) shows the top cell-type-specific genes expressed in the clusters defined in the left panel. **(E)** Force-directed graph drawing of the potentially dividing cells dataset (the left panel), and the violin plot (the right panel) of the expression of cell-type marker genes. MCM2, minichromosome maintenance complex component 2; TOP2A, DNA topoisomerase II alpha; PCNA, proliferating cell nuclear antigen; eSPN, “eccentric” spiny projection neuron. The expression of top cell-type-specific genes of each cluster is shown in the right panel. **(F)** Gene regulatory network analysis of proliferating cells (E). The node size is proportional to its PageRank (PR) centrality. Nodes highlighted in darkorange/cadetblue colors in the network indicate genes in the intersection of three gene sets: 1) top 500 genes with the biggest changes in PR absolute value between STR and CTL; 2) genes with over 0.75 quantile of PR value in each main community (community consisting of over 5 genes) detected in the network; 3) DEGs between STR and CTL. The Nodes with text labels refer to hub genes with maximal biological relevance. **(G)** Bar plots show the GO enrichment results of up- and down-regulated hub genes in (F).

Weighted gene co-expression network analysis and gene ontology (GO) enrichment analysis of bulk RNAseq data implied that PNS dysregulated gene expression networks related to the regulation of three biological functions: transferase activity (*Dusps1, 6, 5, Rgs, Spry2*; the purple module), cellular response to morphogens (*Shisa2, Ccn1, Spry2, Bambi, Smad7*; the purple module), and gliogenesis (*Mag, Trf, Myrf, Plp1, Gjc2*; the magenta module) in adult male offspring (Fig.4B, C). Many genes in the purple module are critical components of the RAS-RAF-MEK-MAPK(ERK) signaling cascade, which is the major downstream of growth factor signaling (e.g., EGF, FGF, VEGF) to promote cell proliferation [14, 18, 45-48]. GO enrichment analysis of the greenyellow and pink modules did not identify any signaling pathways. The major sex differences between males and females were the regulation of the immune effector process and cytokine production (the yellow module) (Fig.4B, C, Sup.Fig.5A) [49].

To further examine PNS-elicited changes in the E-SVZ, we carried out snRNAseq. The nuclei isolated from the E-SVZ cells of different litters were pooled, purified, and subjected to droplet-based snRNAseq (Fig.4A). Quality control filtering and integration of CTL and STR datasets were performed using the Seurat package[50] and the SingCellaR package [51, 52], which generated a merged dataset consisting of 7,407 cells and 20,634 genes for further analysis. According to marker gene expression, we annotated the 18 clusters to 10 different types of cells: 1) *Foxj1*^+^ ependymal cell (EpC, cluster 6); 2) *Prom1*^+^ endothelial-like cell (ElC, cluster 17); 3) dorsal neural progenitor cell (NPCd, cluster 14); 4) neuroblast (NB, clusters 7 and 11); 5) *Drd1*^+^ spiny projection neuron (D1 SPN, clusters 2, 3, 10, 12); 6) *Drd2*^+^ SPN (D2 SPN, clusters 1, 8, 13); 7) ventral neural progenitor cell (NPCv, cluster 16); 8) astrocyte (AC, clusters 4, 5); 9) oligodendrocyte (ODC, cluster 9) and oligodendrocyte precursor cell (OPC, cluster 18); 10) microglial cell (MG, cluster 15) (Fig.4D, Sup.Fig.5C). Cluster 11 NBs are dividing cells expressing *Mki67* and *Top2a*, while cluster 7 NBs are postmitotic cells. Cluster 14 NPCs express *Egfr, Mki67*, and *Btg2*, but not the NB markers *Dcx* and *Tubb3*. Astrocytes are separated into *Aqp4*^-^ (cluster 4) and *Aqp4*^+^ (cluster 5) cells. We detected a significant number of D1 and D2 SPNs that reside close to the E-SVZ (Sup.Fig.5B,C). Compared to the CTL, there were no obvious cell type changes in STR samples (Sup. Fig.4C).

To determine potential intrinsic programs that control NSC/NPC proliferation, we reanalyzed proliferating neural lineage cells. These cells express at least one of the cell cycle markers, *Mki67, Mcm2, Top2a*, and *Pcna* (UMI>0). We identified eight types of cells: EpC (cluster 7), ElC (cluster 8), NPC (cluster 3), NB (clusters 4 and 6), AC (cluster 5), D1 SPN (cluster 1), D2 SPN (cluster 2), and “eccentric” eSPN (cluster 9) (Fig.4E, Sup.Fig.5C) [53, 54]. Cell cycle position estimation analysis (estimating a continuous cell-cycle pseudotime by principal component analysis of cell-cycle genes) showed that most cells in cluster 3 (NPC), cluster 4 (NB), and some cells in cluster 8 (ElC) were actively dividing cells, while cells of other clusters were predicted to be in the G1-phase (Sup.Fig.5D, cell cycle score) [55]. Cells in cluster 3 (NPC, *Egfr*^*+*^, *GFAP*^*-*^) were likely to be the previously described activated NSC (aNSC) and/or transit-amplifying cells (TACs, C cells)[56]. Cluster 5 astrocytes expressed *Gfap* and *Pdgfrb*, but not *Egfr* and *Prom1*, suggesting an astroglial lineage (Sup.Fig.5B). It remains to be determined why a small subset of spiny projection neurons (D1, D2, and eSPN) expresses low levels of cell cycle markers. It seems that in addition to olfactory bulb neurons, this region could produce new striatal neurons [57]. Another possibility is that cells in or close to the E-SVZ could sense some environmental cues, and thus express cell cycle markers.

We next employed the bigSCale2 algorithm to construct gene regulatory networks for the proliferating cells[58] (Fig.4F). Compared to the CTL network, the STR network showed a lower edge-to-node density (gene to gene interaction across functional networks (module)) (CTL = 8.17, STR = 7.63) and higher modularity (gene to gene interaction within a module) (CTL = 0.41, STR = 0.54), suggesting that PNS caused a sparser network with a higher gene expression heterogeneity in the E-SVZ of STR offspring. The analysis implicated two major network modules that regulate cell cycle progression and neural circuit maturation (Fig.4F). GO analysis of differentially expressed hub genes showed that PNS promoted the expression of genes that regulated cell cycle progression (e.g., *Atad2, Top2a, Brd8, Atad5, Smchd1, Ncapg2, Pola1, Ezh2*) and neural stem cell maintenance and differentiation (e.g., *Sox2, Sox5, Nfib*, Zmiz1, *Zeb2*), while it downregulated expression levels of genes modulating synapse organization and membrane potential (e.g., *Slc8a2, Kcnd2, Grm5, Nfia, Ntrk3, Slit3*) (Fig.4F,G). More specifically, PNS enhanced the expression of bromodomain-containing genes *Atad2* (ATPase family AAA domain containing 2) and *Brd8* (bromodomain containing 8) that are involved in ESR1 (estrogen receptor alpha) mediated transcriptional regulation of cell proliferation in cancer cell lines [59-61]. PNS increased expression levels of Wnt pathway modulators *Ezh2, Tcf4, Ephb2*, and *Sox2*, which play important roles in sustaining cell stemness and promoting cell proliferations [62-66]. The Wnt and the estrogen signaling pathways are tightly associated with MAPK3/1 (ERK1/2) signaling to stimulate tumorigenesis [26, 67, 68]. Moreover, MAP4K4 (serine/threonine kinase mitogen-activated protein 4 kinase 4) has been identified as a positive regulator of MAPK (ERK) signaling in cancer cells^68^. In addition, previous studies by our group and others have shown that MAPK3/1 are essential in maintaining the proliferation of adult NSC/NPCs [69-71]. Considering the above transcriptomic analysis, we hypothesized that PNS promoted cell proliferation via the activation of the MAPK (ERK) signaling in the E-SVZ NSC/NPCs of adult male offspring (Sup.Fig.6A).

### Rescue assays reveal that PNS upregulates MAPK3/1 phosphorylation

To test this hypothesis, we first carried out an immunostaining assay to examine phosphorylation levels of MAPK3/1 in the E-SVZ of both CTL and STR mice. The antibody was developed to recognize phosphorylated MAPK3/1 at threonine 202/185 and tyrosine 204/187 loci specifically (ThermoFisher 44-680G). We found that immunostaining of phosphorylated MAPK (p-MAPK) showed a punctate pattern (Figs.5A, 6B). This pattern was affirmed by using another p-MAPK3/1 specific antibody from a different vendor (Cell Signaling, 4370) and an antibody against unphosphorylated MAPK(ERK) (Cell Signaling, 4695) (Sup.Fig.6B, C). Furthermore, when treating the section with lambda protein phosphatase (Lambda PP) before primary antibody incubation, the immunoreactivity of the p-MAPK3/1 antibody was completely abolished, suggesting the high specificity of the antibody in recognizing p-MAPK (Sup.Fig.6D).

Since both phosphorylated and unphosphorylated MAKP3/1 staining displayed punctate patterns, we reasoned that the number of puncta correlates with the status of phosphorylation. By employing this quantitative methodology, we next examined the phosphorylation of MAPK3/1 in the E-SVZ at different postnatal developmental time points (P0.5-180). Concomitant with the cell proliferation finding (Fig.1B), we found phosphorylation level of MAPK3/1 was considerably upregulated in the E-SVZ of stressed adult offspring (P60, P180) but not at earlier stages (Fig.5).

**Fig. 5.**
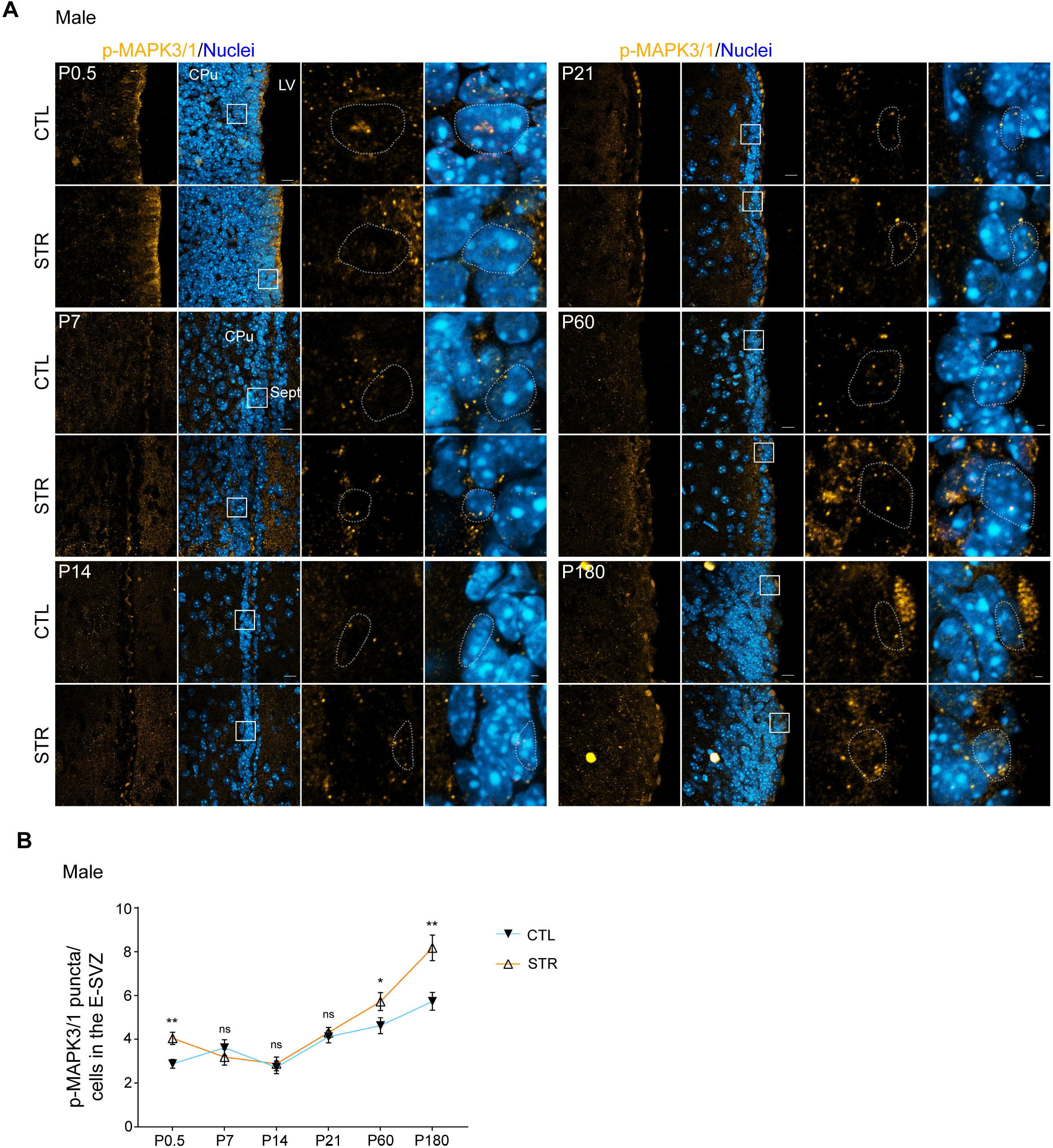
MAPK3/1 phosphorylation in developing and adult brains. MAPK3/1 phosphorylation showed a dramatic difference between CTL and STR at P180 and P60, but not in samples collected from P0.5 to P21. At each time point, the panels on the right showed an enlarged area labeled in the left panels. Scale bars, 10 μm, 1 μm (zoomed-in images). Mann-Whitney test. All data were presented as mean ± SEM. *p < 0.05, **p < 0.01.

To definitively verify the role of the MAPK3/1 pathway in regulating cell proliferation in stressed male mice, we carried out rescue assays by treating the STR and CTL animals with a selective phosphorylated MAPK3/1 inhibitor, LY3214996, and an inhibitor of upstream protein kinases MAP2K1/2 (MEK1/2), GSK1120212. We found that administration of these inhibitors effectively reduced the numbers of p-MAPK3/1 puncta and proliferating cells in the E-SVZ of both CTL and STR mice (Fig.6). Taken together, the rescue assays suggested that MAPK(ERK) signaling played a critical role in promoting NSC/NPC proliferation in STR adult male mice.

**Fig. 6.**
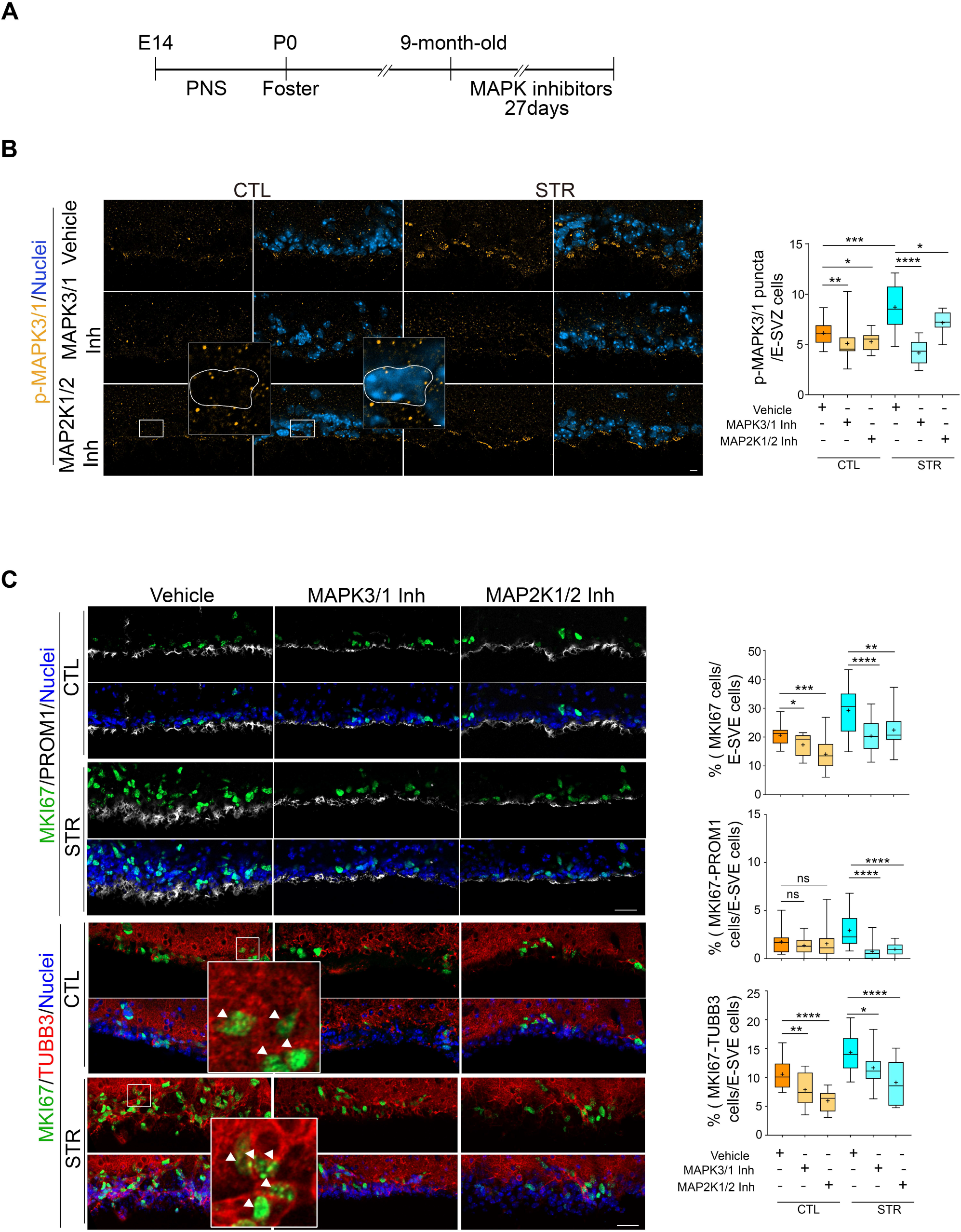
Inhibition of MAPK activity counteracts PNS-triggered NSC/NPC proliferation in the lateral E-SVZ. **(A, B)** Administration of MAPK3/1 inhibitor LY3214996 (LY) and MAP2K1/2 inhibitor GSK1120212 (GSK) decreases the total phosphorylated MAPK3/1 puncta in the E-SVZ (CTL = 6.2 ± 0.3, n = 18; CTL/LY = 5.1 ± 0.5, n = 16; CTL/GSK = 5.3 ± 0.2, n = 17; STR = 8.7 ± 0.5, n = 16; STR/LY = 4.2 ± 0.3, n = 16; STR/GSK = 7.2 ± 0.3, n = 15. CTL vs LY, P = 0.0056; CTL vs GSK, P = 0.0032; CTL vs STR, P = 0.0002; STR vs LY, P < 0.0001; STR vs GSK, P = 0.0405). Scale bars, 30 μm. **(C)** Inhibition of MAPK pathway reduces the number of both PROM1^+^/MKI67^+^ ependymal cells and TUBB3^+^/MKI67^+^ neuroblasts. MKI67^+^cells: CTL = 20.7 ± 0.8%, n = 18; CTL/LY = 17.3 ± 1.0%, n = 15; CTL/GSK = 14.1 ± 1.6%, n = 12; STR = 29.2 ± 1.7%, n = 24; STR/LY = 20.3 ± 0.8%, n = 38; STR/GSK = 22.4 ± 0.9%, n = 38. CTL vs LY, P = 0.0229; CTL vs GSK, P = 0.0008; STR vs LY, P < 0.0001; STR vs GSK, P = 0.0013. PROM1^+^/MKI67^+^ cells: CTL = 1.8 ± 0.3%, n = 16; CTL/LY = 1.4 ± 0.2%, n = 15; CTL/GSK = 1.4 ± 0.5%, n = 12; STR = 2.9 ± 0.4%, n = 20; STR/LY = 0.8 ± 0.2%, n = 17; STR/GSK = 1.0 ± 0.2%, n = 14. CTL vs LY, P = 0.5263; CTL vs GSK, P = 0.3770; STR vs LY, P < 0.0001; STR vs GSK, P < 0.0001. TUBB3^+^/MKI67^+^ cells: CTL = 10.6 ± 0.6%, n = 18; CTL/LY = 7.9 ± 0.7%, n = 15; CTL/GSK = 6.0 ± 0.5%, n = 12; STR = 14.3 ± 0.8%, n = 20; STR/LY = 11.7 ± 0.7%, n = 17; STR/GSK = 9.1 ± 1.0%, n = 14. CTL vs LY, P = 0.0091; CTL vs GSK, P < 0.0001; STR vs LY, P < 0.0151; STR vs GSK, P < 0.0010. Scale bars, 30 μm. Mann-Whitney test. All data were presented as mean ± SEM. *p < 0.05, **p < 0.01, ***p < 0.001, ****p < 0.0001.

## Discussion

Psychosocial, cultural, and environmental stressors experienced during gestation can be detrimental to pregnancy as well as maternal and fetal health. Recent studies have suggested that prenatal maternal stress can shape neural circuitry, neural cell fate, and behavior in offspring, which may set the stage for postnatal disease vulnerability [7, 9-11]. Neural stem cells in the adult central nervous system are highly sensitive to environmental stimuli. For example, neural stem cells can rapidly respond to brain and spinal cord injuries and mitogen stimulation (e.g., EGF, VEGF, and FGF) to proliferate [14, 18, 72, 73]. Moreover, under normal physiological conditions in rodents, growth factors EGF and VEGF have been shown to activate dormant/quiescent proliferation [14, 72]. In the current study, we showed the long-lasting response of neural stem cells in STR offspring to a prenatal adverse stimulus, restraint stress. Our data suggested that in adult STR offspring, cells in the stem cell niche E-SVZ responded to the stress through enhancing growth factors-, Wnt-, and estrogen-regulated pathways, and the MAPK (ERK) signaling cascade could be the central player that mediates the long-lasting effect of the prenatal event.

It has now been accepted that adult neurogenesis happens in mammals despite some disputes over the origin of neural stem cells in rodent brains and the existence of neurogenesis in humans [14, 15, 18, 48, 71, 74-77]. In mammals, the current theory of adult neurogenesis is adopted from the well-established embryonic neurogenesis paradigm that neurons and glial cells are generated from sequential symmetrical or asymmetrical divisions of neuroepithelial cells/radial glia and then intermediate progenitors [78-80]. Our snRNAseq analysis suggested that PNS upregulated the expression level of *Zeb2* (zinc finger E-box binding homeobox 2) and its downstream target *Sox6* in the germinal zone of adult STR brains [81] (Fig.4F). ZEB2 is one of the essential transcription factors that induce epithelial-mesenchymal transition (EMT) during pluripotent stem cell differentiation, embryonic development, wound healing, and tumorigenesis [82-84]. Studies by Miquelajauregui et al. and Deryckere et al. found that ZEB2 plays a critical role in controlling neurogenesis in prenatal and postnatal brains, respectively [81, 85]. Moreover, Wnt and MAPK (ERK) pathways have been shown to induce *Zeb2* expression and promote EMT in cancer cells [82, 83]. MAPK(ERK) is a potent signaling transducer of EGF and VEGF, which are essential for promoting dormant or quiescent neural stem cell activation and neural progenitor cell proliferation [14, 19, 24, 46, 70-72]. Therefore, PNS could activate dormant/quiescent neural stem cell proliferation via promoting MAPK (ERK)-mediated EMT in the germinal zones of the adult brain.

To examine the possible association between PNS and basic olfactory function, we carried out the buried food test and the habituation/dishabituation test. Our initial study showed that STR male mice (5-8-month-old) had a largely intact olfactory function to locate food pellets and discriminate nonsocial odors, while the STR mice showed impairment in the differentiation of different social odors and habituation of social odor 2 (STR Social 1-3 median=2.98 sec to Social 2-1 median=2.58 sec; CTL Social 1-3 median=1.34 sec to Social 2-1 median=4.66 sec) (Sup.Fig.4B-D). Calbindin 2 (CALB2, a.k.a. Calretinin) staining showed a disorganized glomerular layer (Gl) and a slightly increased number of CALB2^+^ cells in the external plexiform layer (EPl) and the granule cell layer (GrO) of the OB in stressed male mice (Sup.Fig.4A). Glomeruli and granular cells have been shown to play essential roles in the regulation of olfactory habituation behavior [86, 87]. Moreover, scRNAseq analysis showed that PNS attenuated the expression of genes modulating synapse organization and membrane potential, which may affect olfactory functions (Fig.4G). For example, PNS downregulated the expression of glutamate metabotropic Receptor 5 (*Grm5*, a.k.a. mGluR5), which has been shown to regulate granule cell-mediated inhibition and odor discrimination [88, 89]. Taken together, although the causal relationship between increased NSC/NPC proliferation and the behavioral deficit is still elusive, BrdU lineage tracing together with immunostaining showed disorganized cytoarchitecture of the olfactory glomerular layer and slightly increased numbers of CALB2+ cells in the external plexiform layer and the granule cell layer, which have been shown to play essential roles in the regulation of olfactory habituation behavior [90-93].

## Materials and Methods

### Experimental Animals

Mice were kept and fed in standard conditions on a 12hr light/dark cycle. Experimental procedures on animals were performed following the guidelines of the University of California, Los Angeles (UCLA) Institutional Animal Care and Use Committee and UCLA Animal Research Committee, and the Regional Committee for Medical Research Ethics of Tongji University. The inbred strain C57BL/6NCrlVr mice were used in this study. Pregnancy was timed by daily monitoring for vaginal plugs.

### Restraint stress and tissue collection

To induce stress between gestational days 14 and delivery (∼E19.5), the pregnant dam was placed individually in a plastic restrainer fitted closely to body size for 60 min/day (session at 0800h). Control pregnant females were left undisturbed in their home cages. After delivery, the prenatally stressed (STR) and the CTL pups were fostered by unstressed, same strain mothers at P0. All pups were weaned at P21. For tissue collection, the animals were sacrificed, and their brains were removed for immunostaining or RNA sequencing.

### Buried food and olfactory habituation/dishabituation tests

The tests were described by Yang and Crawley[94]. Details are given in SI Appendix.

### Bromodeoxyuridine (BrdU) administration

The BrdU injection was described in our previous study[50]. Details are given in SI Appendix.

### Immunohistochemistry

The primary antibodies used in this study were BrdU (Abcam ab6326, 1:100; ab1893 1:250), MKI67 (Abcam ab15580, 1:200), GFAP (ThermoFisher PA1-10004, 1:500), TUBB3 (R&D MAB1195, 1:500), CALB2 (Synaptic Systems 214111, 1:200), DCX (Santa Cruz sc-8066, 1:400), RBFOX3 (Abcam ab177487, 1:500), p-MAPK3/1(ThermoFisher 44-680G, 1:100), p-MAPK3/1 (Cell Signaling 4370, 1:200), and MAPK3/1 (Cell Signaling 4695, 1:250). Fluorophore-conjugated secondary antibodies were purchased from ThermoFisher Scientific and Jackson ImmunoResearch.

### Terminal deoxynucleotidyl transferase dUTP Nick-End Labeling (TUNEL) assay

TUNEL assay was carried out following the manufacturer’s instructions (*In Situ* Cell Death Detection Kit, Roche-11684795910).

### Tissue processing for BrdU immunostaining

Details are given in SI Appendix.

### Transcriptomic analysis

The brains of 7- or 8-month-old STR and CTL offspring were sectioned at 300μm thickness in artificial cerebrospinal fluid supplied with actinomysin D (3μM, Sigma-Aldrich). The lateral walls of ependymal SVZ (E-SVZ) were dissected, frozen in liquid nitrogen, and stored in air-tight tubes in -80°C freezer.

#### Bulk RNAseq

Total RNA was extracted and purified using RNeasy Mini Kit (Qiagen 74104). 100ng of qualified RNA was subjected to VAHTS™ mRNA-seq V3 Library Prep Kit for Illumina^®^ (Vazyme NR611) for library preparation, followed by quality assessment using Agilent Bioanalyzer 2100, and sequencing on Illumina NovaSeq 6,000. The mRNA sequencing reads were mapped to the mouse reference genome (GRCm38) using HISAT2 (version 2.2.1) and StringTie (version 2.1.4), and the mapped reads were assigned using featureCounts function of R package Rsubread (version 2.0.1). The counts were normalized using the DESeq2 R package (version 1.30.0)[95]. Pseudogenes (gene names with the prefix “Gm”) were filtered out, and top 10,000 highly variable genes were kept for weighted gene co-expression network analysis (WGCNA) using the R package WGCNA (version 1.69)[96]. Specifically, a soft-power of 12 was chosen to construct a topological overlap matrix from the gene correlation network. Modules were determined by the Dynamic Hybrid Cut algorithm using a “deepSplit” parameter of 2. Highly correlated modules (Pearson correlation of module eigengene > 0.9) were merged as one module using a “mergeCutHeight” parameter of 0.25. Gene ontology (GO) and KEGG pathway enrichment analyses of genes in each identified module were performed using the R package clusterProfiler (version 4.4.4)[97] set at default parameters.

#### Single nucleus RNAseq (snRNAseq)

Single nuclei were extracted from pooled frozen lateral walls of ependymal SVZ (E-SVZ) from CTL and STR mice using a previously described ultracentrifugation method[98] with some modifications. The single nuclei RNA sequencing library construction was performed using the Chromium Next GEM Single Cell 3’ Reagent Kits v3.1 (10× Genomics) according to the user guide. The quality assessment of the libraries was conducted using Agilent Bioanalyzer 2100, and the sequencing was performed on an Illumina Novaseq 6000. The alignment, filtering, barcode assignment, and UMI counting of the sequenced reads were processed using the Cell Ranger pipeline (version 5.0.1, 10× Genomics). Quality control (QC) of cells and genes was performed using R package Seurat (version 4.1.1). Doublets were removed using R package scDblFinder (version 1.10.0). Additional downstream workflow (including further QC, sample integration, highly variable gene selection, dimensional reduction, clustering, cluster marker gene identification, and visualization) was processed via SingCellaR R package (version 1.2.1)[51, 52]. According to the expression of the marker genes, along with well-established celltype specific gene sets, the 18 clusters were assigned to 11 major celltypes: dorsal neural progenitor cells (NPCd), ventral neural progenitor cells (NPCv), neuroblasts (NB), astrocytes (AC), ependymal cells (EpC), endothelia-like cells (ElC), *Drd1*^+^ SPN (D1), *Drd2*^+^ SPN (D2), oligodendrocyte progenitor cells (OPC), oligodendrocytes (ODC), and microglia (MG).

Cells (except for clusters without neuronal lineage potential, OPC, ODC and MG) expressing at least one of the four cell cycle-related genes (*Mki67, Top2a, Mcm2*, and *Pcna*) were isolated and re-clustered via SingCellaR workflow. The potentially dividing cells were clustered into 9 clusters and annotated as neural progenitor cells (NPC), AC, NB, EpC, ElC, D1, D2, “eccentric” spiny projection neuron (eSPN), respectively. The cell-cycle position of each potentially dividing cell was estimated using the tricycle R package (version 1.4.0)[55]. The putative gene-to-gene correlated network of the potential dividing cells was inferred using the bigSCale2 algorithm (version 2.0)[58]. The network centrality PageRank (PR) was chosen to represent gene essentiality. Gene communities in each network were identified, and those communities consisting of over 5 genes were considered as main communities. Hub genes with maximal biological relevance in each main community were defined as the intersection of three gene sets: 1) top 500 genes with the biggest change in PR absolute value between STR and CTL; 2) genes with over 0.75 quantile of PR value; 3) DEGs between STR and CTL determined using SingCellaR package (version 1.2.1)[51, 52]. GO enrichment analysis of the hub genes was performed using R package clusterProfiler (version 4.4.4)[97] set at default parameters. Details are given in SI Appendix.

### Rescue assay

A MAPK3/1 inhibitor LY3214996 (Selleck, 16 mg/kg) and a MAP2K1/2 inhibitor GSK1120212 (Selleck, 0.35 mg/kg), and the vehicle solution were daily intraperitoneally (IP) injected to 9-to 10-month-old prenatally stressed male offspring for 27 days (except for the 11^th^ and 22^nd^ days), respectively. Details are given in SI Appendix.

### Lambda Protein Phosphatase (Lambda PP) reaction

The assay was carried out following the manufacturer’s instructions (NEB, P0753S). Details are given in SI Appendix.

### Imaging and Statistical Analysis

At least three mice of each gender from different litters at different postnatal stages were randomly selected for immunostaining and quantification analysis. Stained sections were imaged with Zeiss LSM800 confocal microscope as tiled or single images. Images obtained were processed and quantified with Imaris software and ImageJ. Statistical analyses of the data were carried out with R-package, GraphPad Prism, and SAS (Statistical Analysis Software). Mann-Whitney test was used to assess statistical significance between independent experimental groups. All reported significant levels to represent two-tailed values. Details are given in SI Appendix.

## Acknowledgments

The study was supported by the National Key Research and Development Program of China (2020YFC2002804), NIH Grants (5R21MH115382, U54HD087101, and P50HD103557), State Key Program of the National Natural Science Foundation of China (82030035), the Foundation of Shanghai Municipal Education Commission (2019-01-07-00-07-E00055), Peak Disciplines (Type IV) of Institutions of Higher Learning in Shanghai, Postdoctoral Science Foundation funded project (2022M712418), the 2021 Shanghai “Super Postdoctoral” Incentive Program, Key R&D Program of Jiangsu (BE2020026), and National Natural Science Foundation of China (31872761). We thank the Intellectual and Developmental Disabilities Research Center at the University of California, Los Angeles.

---

**Sup. Figure 1.**
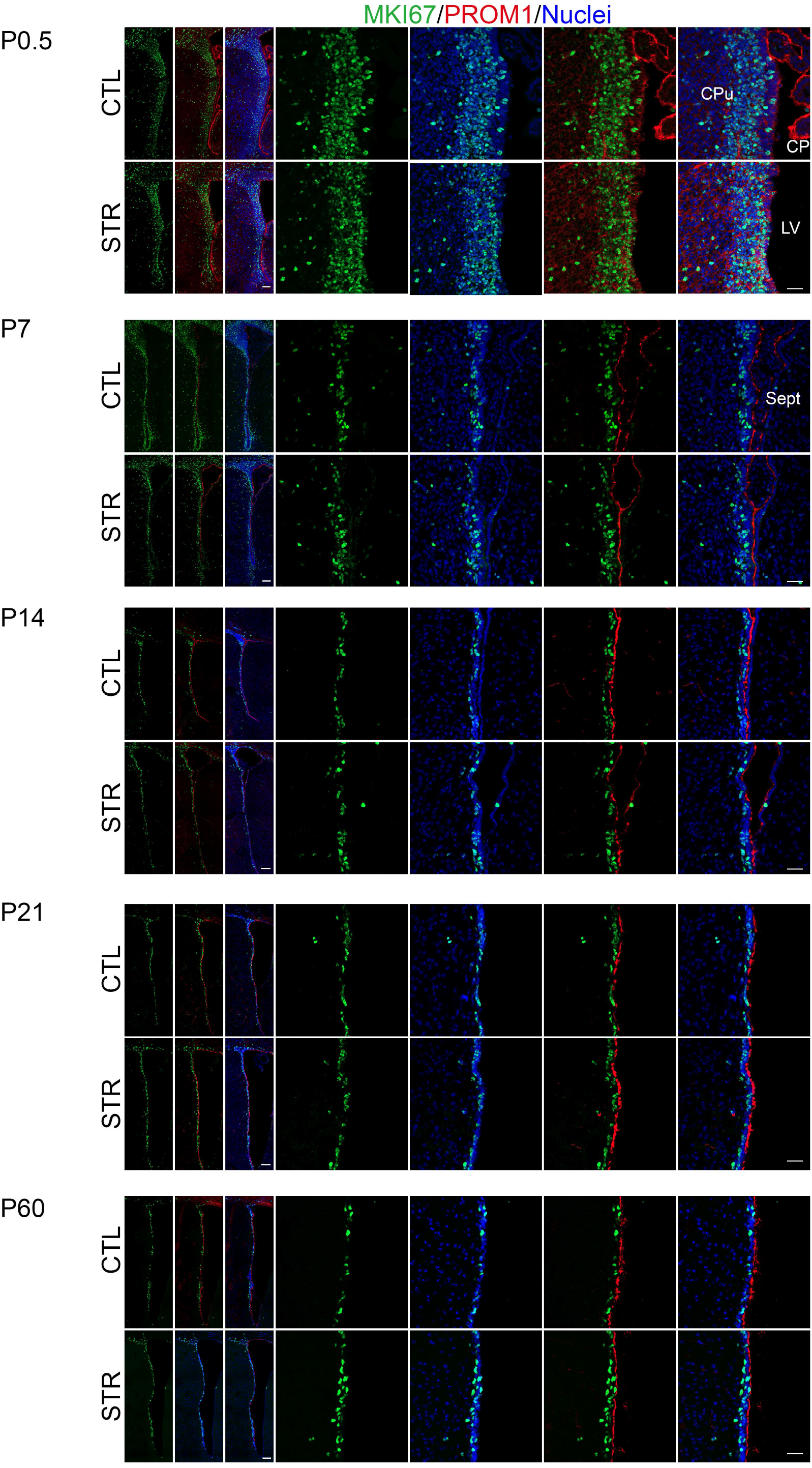

**Sup. Figure 2.**
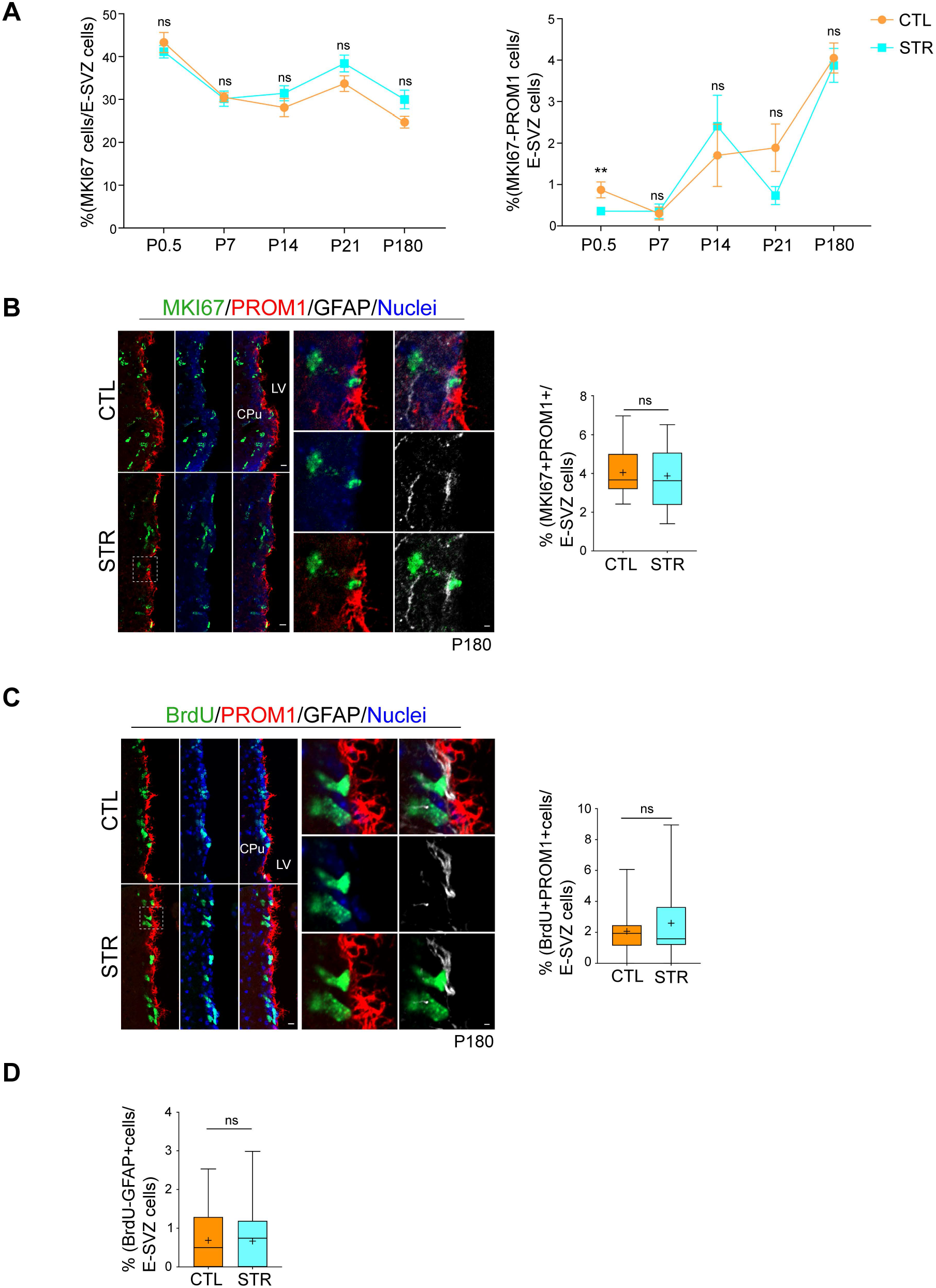

**Sup. Figure 3.**
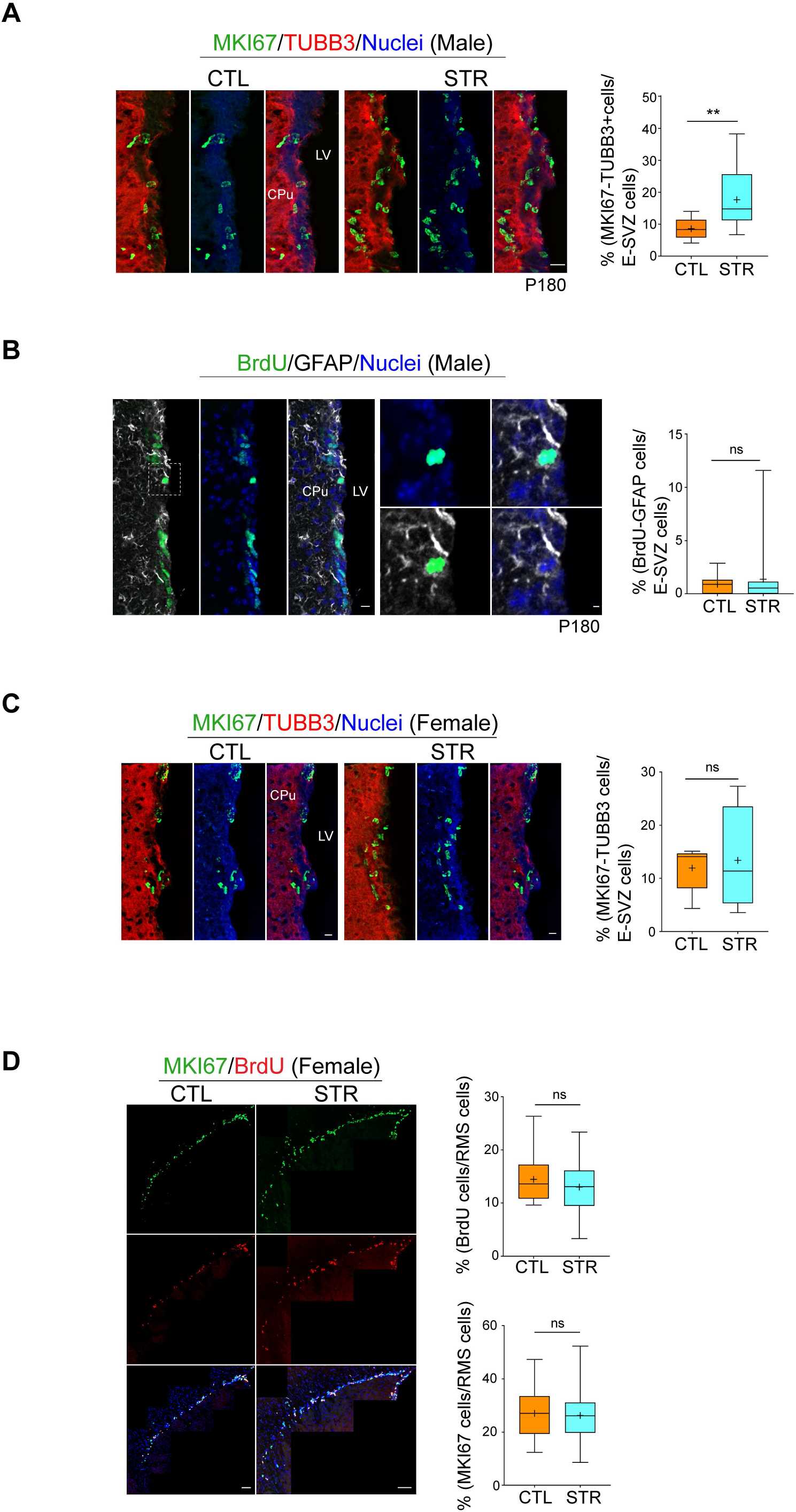

**Sup. Figure 4.**
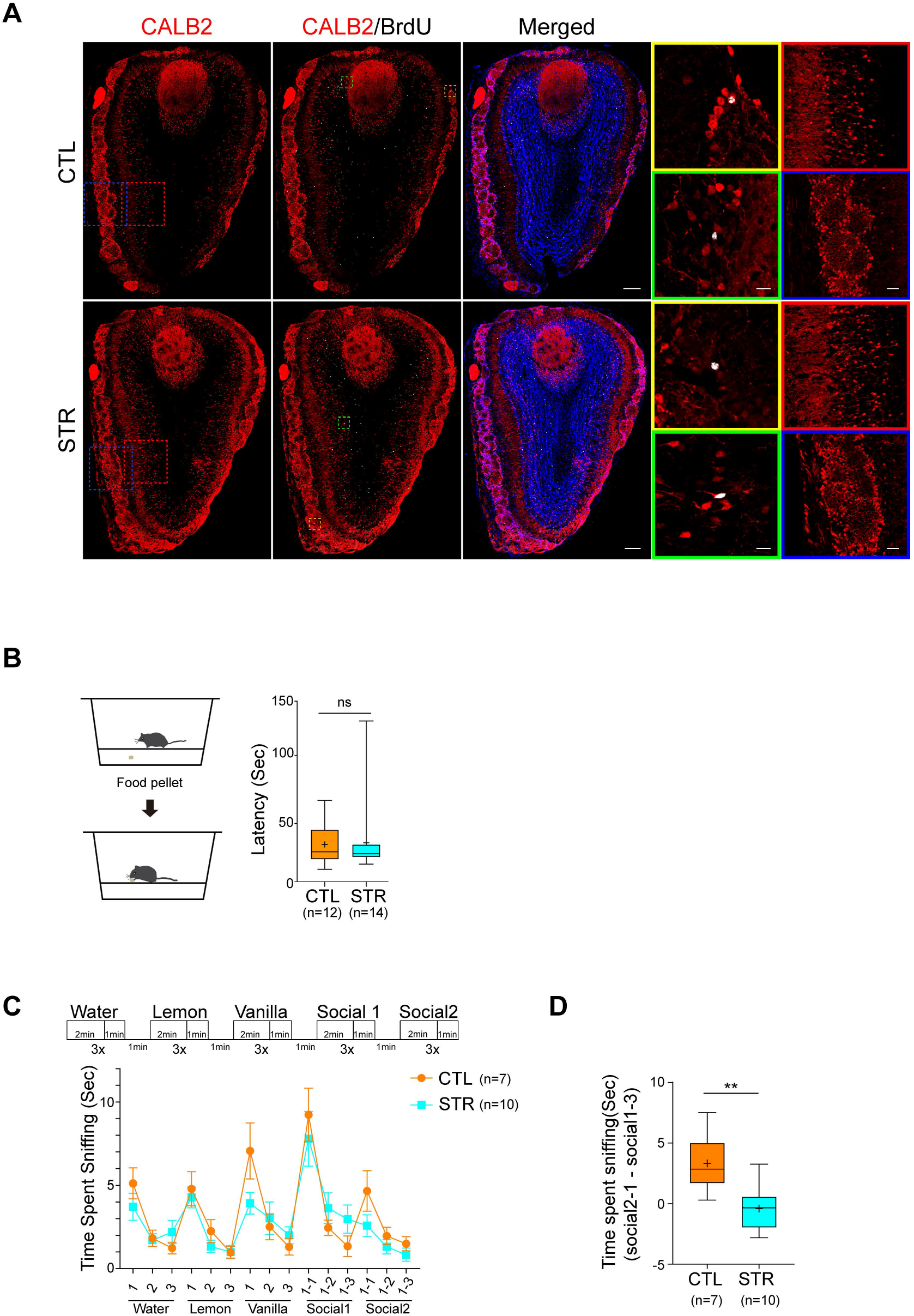

**Sup. Figure 5.**
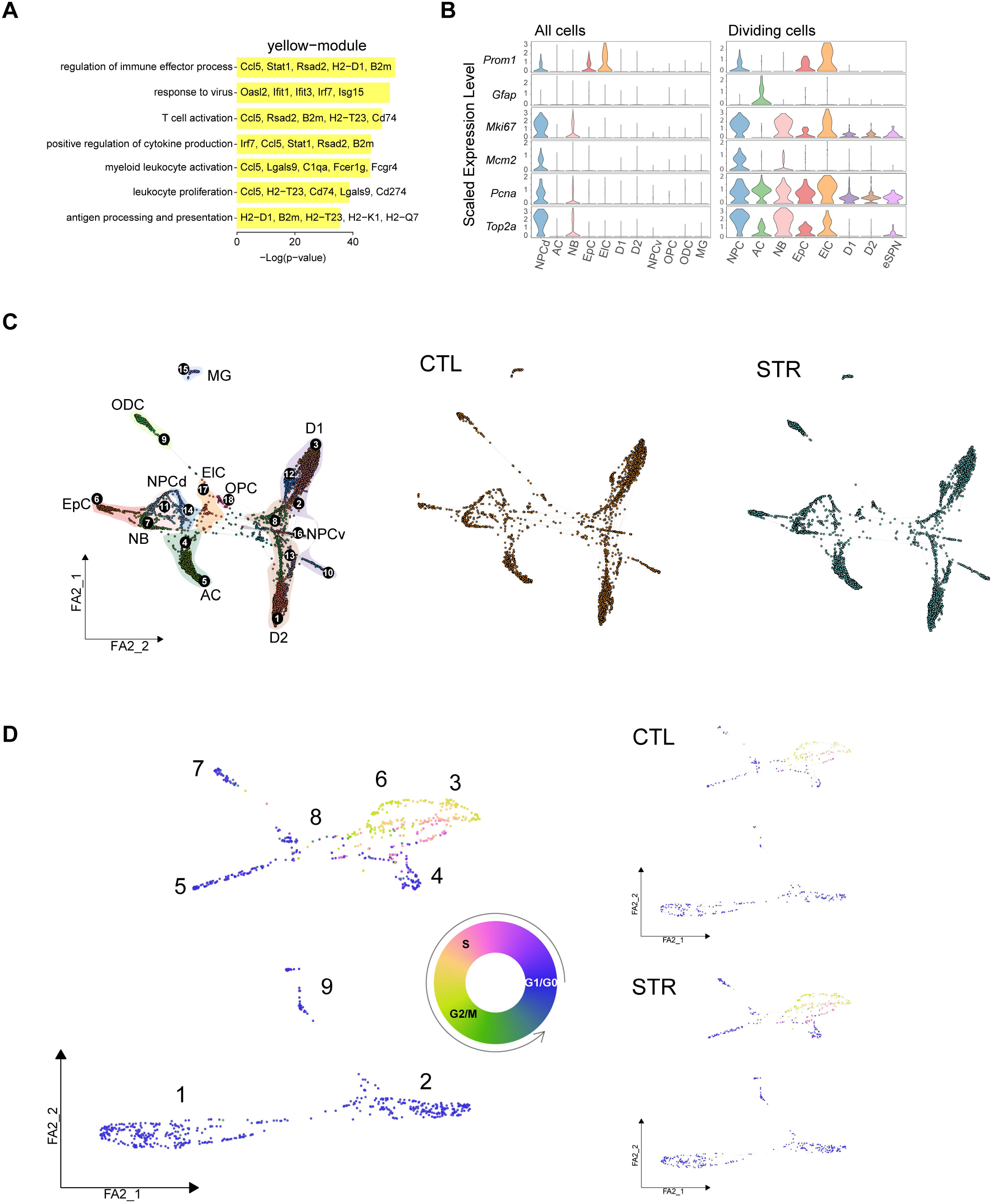

**Sup. Figure 6.**
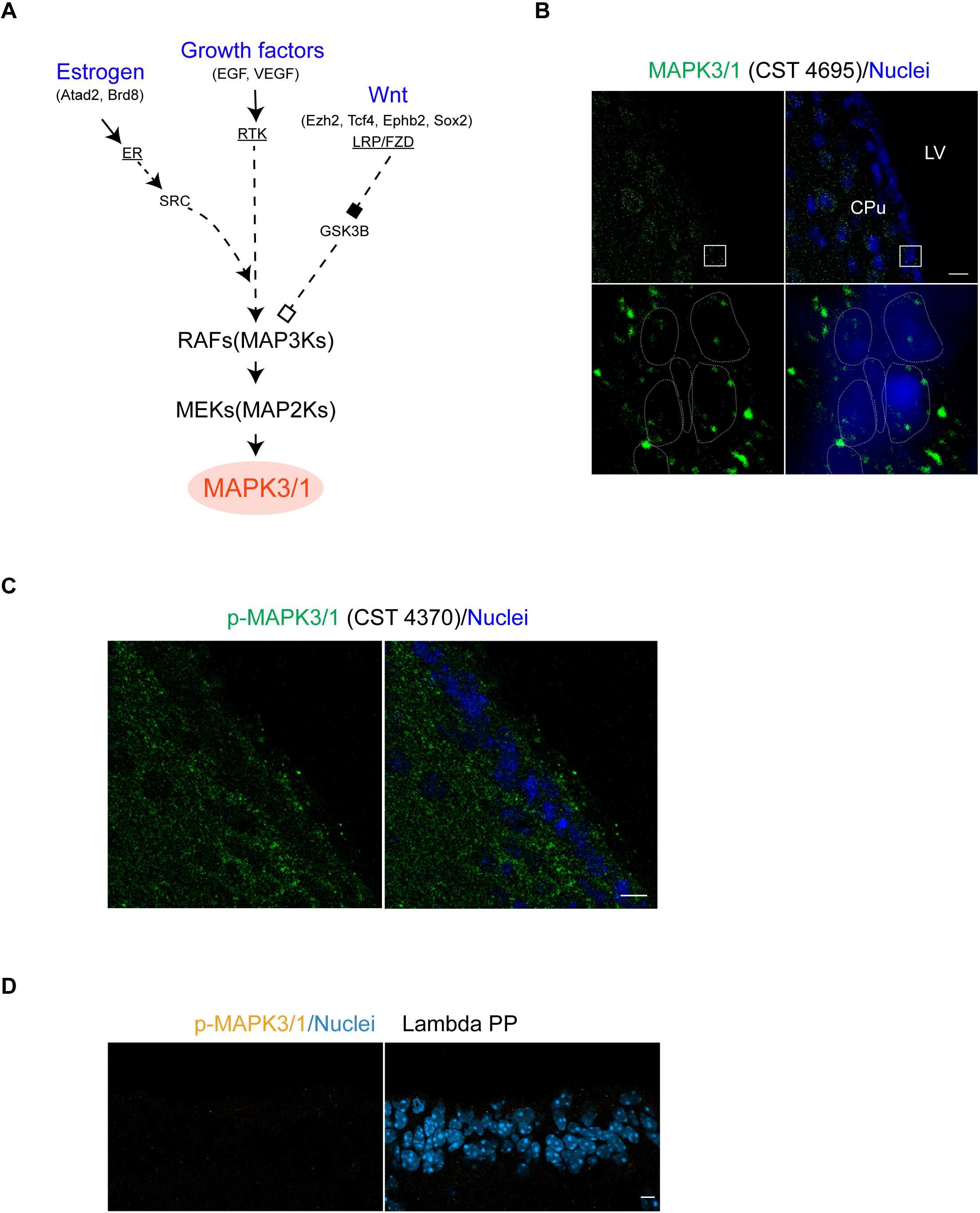

**Sup. Table 1.**
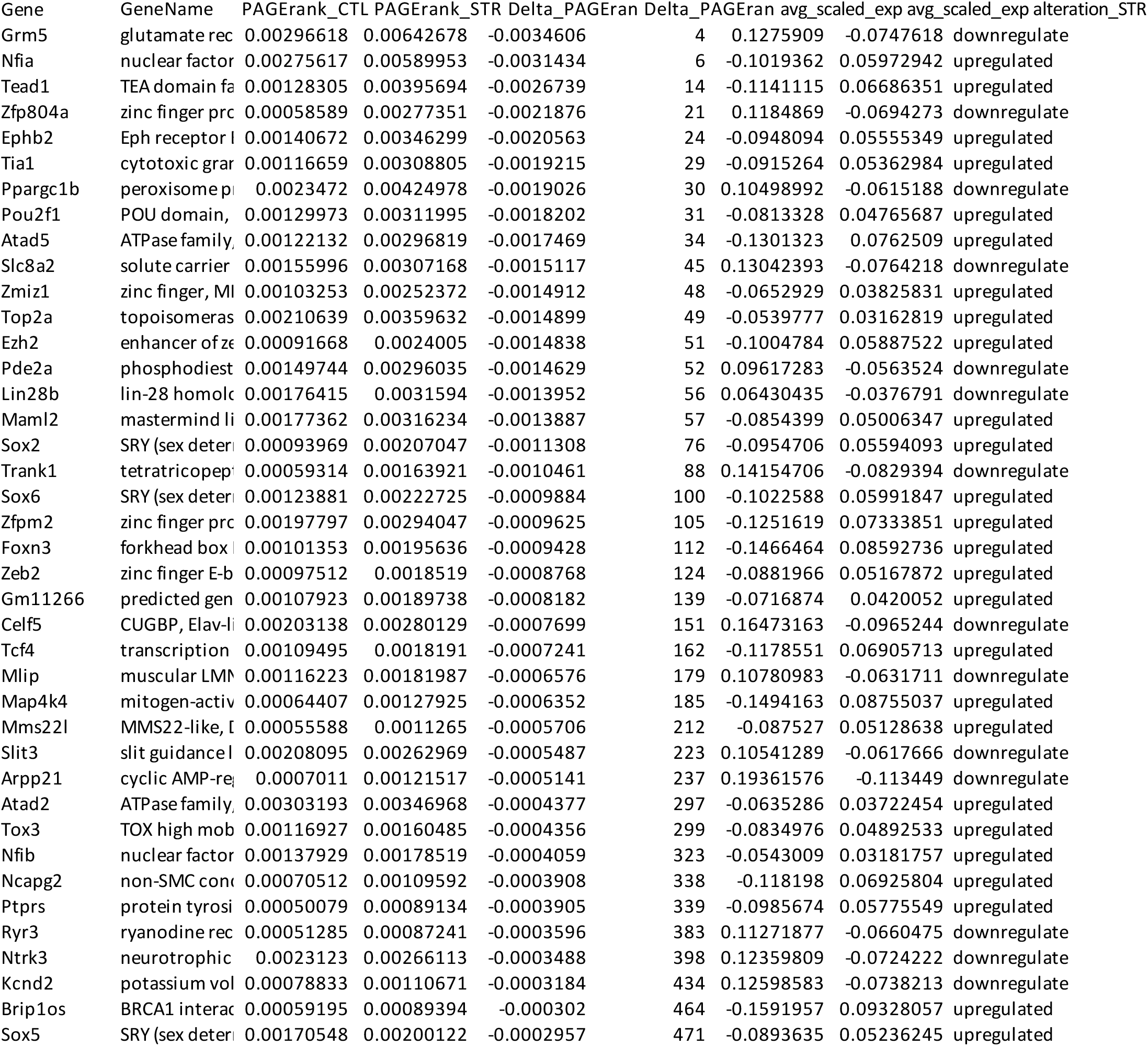

