## Supplementary material for "Transcriptomic analyses reveal prenatal stress promotes late-onset neural stem cell proliferation in adult male offspring": Sup info

**Materials and Methods**

**Experimental Animals.** Mice were kept and fed in standard conditions on a 12-hr light/dark cycle. Experimental procedures on animals were performed in accordance with the guidelines of UCLA Institutional Animal Care and Use Committee and UCLA Animal Research Committee, and the Regional Committee for Medical Research Ethics of Tongji University. The inbred strain C57BL∕6NCrlVr mice were used in this study. Pregnancy was timed by daily monitoring for vaginal plugs.

**Restraint stress and tissue collection.** To induce stress between gestational days 14 and delivery (~E19.5), the pregnant dam was placed individually in a plastic restrainer (Stoelting 51332, Fisher Scientific, or homemade 3D-printed restrainer) fitted closely to body size for 60 min/day (session at 0800h). Control pregnant females were left undisturbed in their home cages. To eliminate the postnatal stress effects from their own stressed mother, the STR pups were fostered by unstressed, same strain mother at P0. To eliminate the possible effects of fostering, the control offspring were fostered by unstressed same strain mother at P0. At 21d, females and males were separated from their mothers and group housed with same-sex littermates until adulthood. For immunostaining related assays, mice were overdosed with isoflurane and perfused intracardially with PBS followed by 4% paraformaldehyde (PFA), and the brains were removed at different postnatal time points (i.e., P0.5, P7, P14, P21, P60, P180, 8-9 months, 10-11 months).

For RNA sequencing analyses, mice (postnatal ~ 8 months, 15 females and 10 males in the control fostered group (CTL), and 10 females and 14 males in the prenatally stressed group (STR)) were transcardiac perfused with saline, and brains were immediately removed and submerged in fresh ice-cold ACSF (124 mM NaCl (Sigma-Aldrich, S3014), 2.5 mM KCl (Sigma-Aldrich, P9333), 1.2 mM NaH_2_PO_4_ (Sigma-Aldrich, S3139), 24 mM NaHCO_3_ (Sigma-Aldrich, S5761), 5 mM HEPES (Sigma-Aldrich, S3375), 13 mM glucose (Sigma-Aldrich, G8270), 2 mM MgSO_4_ (Sigma-Aldrich, 746452), and 2 mM CaCl_2_ (Sigma-Aldrich, C5670) with a pH of 7.3 - 7.4) pre-bubbled with carbogen (95% oxygen and 5% carbon dioxide) adapted from a previously described formula[1]. The brains were then sectioned into 300 µm slices using a vibratome (Leica VT1200S) situated on ice. Each slice was immediately transferred to a Petri dish (60 mm in diameter) containing pre-carbonated ice-cold NMDG-ACSF (93 mM NMDG (Sigma-Aldrich, M2004), 2.5 mM KCl, 1.2 mM NaH_2_PO_4_, 30 mM NaHCO_3_, 20 mM HEPES, 25 mM glucose, 10 mM MgSO_4_, 0.5 mM CaCl_2_, 5 mM sodium L-ascorbate (Sigma-Aldrich, A4034), 2 mM thiourea (Sigma-Aldrich, T7875), 3 mM sodium pyruvate (Sigma-Aldrich, P2256) with a pH of 7.3 - 7.4, and 30 µM ActD (Sigma-Aldrich, A1410)). Bilateral walls of the lateral ventricles containing the ependymal-subventricular zone were microdissected from the slices using a microsurgical stab knife, and immediately transferred to 1.5-mL cryovials. The vials were snap-frozen in liquid nitrogen for ~ 2 min, and then stored at -80 ℃ until use.

The sample size was determined based on our previous experience and a recently published study [2].

**Behavior tests.**

Buried food test. The test was described by Yang and Crawley [3]. In brief, 24-hr before the test, all chow pellets were removed from the food hopper of the home cage. In each trial, a single mouse was placed at random in a test cage to recover a 1.5 g food pellet buried approximately 1 cm below the surface of a 3 cm deep layer of mouse bedding material. The location of the food pellet was changed in a random fashion. The latency to find the food pellet was defined as the time between when the mouse was placed in the cage and when the mouse uncovered the food pellet and grasped it in its forepaws and/or teeth.

Olfactory habituation/dishabituation test.

The test was described by Yang and Crawley [3]. In brief, the exploration sequence consists of water, two nonsocial odors (orange and vanilla extract), and two social odors. Each odor (or water) is presented in three consecutive trials of 2 min per trial. The inter-trial interval is 1 min. The different stimuli were presented in the same order for each mouse.

All mice tested were about 8-9-month-old. Behavior was video-recorded and analyzed offline. The time spent investigating the stimulus was measured using a stopwatch.

**Bromodeoxyuridine (BrdU) administration.** BrdU (Sigma, B5002) was suspended in water to a working concentration of 10mg/ml. Prior to injection, animals were weighed. After cleaning the injection site with ethanol, BrdU solution was administered through IP injection at 200mg/kg (adult mice).

**Terminal deoxynucleotidyl transferase dUTP Nick-End Labeling (TUNEL) assay.** TUNEL assay was carried out following manufacturer’s instructions (*In Situ* Cell Death Detection Kit, Roche-11684795910). For a positive control, sections were incubated for 15 min at 37 °C in proteinase K (20 μg/ml), followed by treatment with DNase I (3000 U/ml) for 10 min at room temperature.

**Tissue processing for BrdU immunostaining.** For BrdU/CD133/GFAP staining, the sections were pretreated with 2M HCl for 20 minutes at 37°C, and then neutralized with 0.1M sodium borate buffer (pH8.5) prior to blocking. For BrdU/MKI67/DCX staining, the sections were pretreated with 10mM citric acid (pH 6.0) for 15 minutes in a steamer at ~99°C during the entire tissue incubation time, then cool slides for 2min at room temperature covering with citrate buffer.

**Transcriptomic analysis.** The brains of 7- or 8-month-old STR and CTL offspring were sectioned at 300 μm thickness in artificial cerebrospinal fluid supplied with actinomysin D (3μM, Sigma-Aldrich). The lateral walls of ependymal SVZ (E-SVZ) were dissected, and snap frozen in liquid nitrogen, and stored in air-tight tubes in -80°C freezer.

**Bulk RNAseq and transcriptomic analysis.**

Total RNA of the CTL and STR mice were extracted from frozen tissue and purified using the RNeasy Mini Kit (QIGEN 74104) according to manufacturer’s instructions. 100 ng of qualified RNA was isolated for library preparation using the VAHTSTM mRNA-seq V3 Library Prep Kit for Illumina (Vazyme NR611). The quality of the library preparation was assessed using the Agilent Bioanalyzer 2100, and the total RNA was sequenced using the Illumina NovaSeq 6000.

The mRNA sequencing data were mapped to the mouse genome (GENCODE (*https://www.gencodegenes.org/mouse/*) M25, GRCm38.p6) using HISAT2 (version 2.2.1) and StringTie (version 2.1.4)[4], and the mapped reads were assigned using the “featureCounts” function of the R package Rsubread (version 2.0.1). The counts were normalized using the DESeq2 R package (version 1.30.0)[5]. Pseudogenes (gene names with the prefix “Gm”) were filtered out, and top 10,000 highly variable genes were kept for weighted gene co-expression network analysis (WGCNA) using the R package WGCNA (version 1.69) [6]. Specifically, a soft-power of 12 was chosen to construct a topological overlap matrix from the gene correlation network. Modules were determined by the Dynamic Hybrid Cut algorithm using a “deepSplit” parameter of 2. Highly correlated modules (Pearson correlation of module eigengene > 0.9) were merged as one module using a “mergeCutHeight” parameter of 0.25. Gene ontology (GO) and KEGG pathway enrichment analyses of genes in each identified module were performed using the R package clusterProfiler (version 4.4.4) [7] set at default parameters.

**Single nucleus RNAseq(snRNAseq) and data analysis.**

To isolate and purify the nuclei, frozen tissue from CTL and STR mice were collected and pooled into 4 samples (CTL-1, CTL-2, STR-1, STR-2) where each sample composed of 3 independent microdissected laterals wall from 3 independent mice for single nuclei sequencing. Nuclei were isolated and purified as previously described[8] with some modifications. In brief, 4.41 mL of sucrose cushion buffer (1.8 M sucrose (Sigma-Aldrich, V900116), 10 mM Tris-HCl pH 8.0 (ThermoFisher Scientific, 15568025), 3 mM MgAc_2_ (Sigma-Aldrich, M5661), 3 µM ActD, 1 tablet of protease inhibitor cocktail (Sigma-Aldrich, 1169749800)) was added to the bottom of an ultracentrifuge tube (13.2 mL, Beckman, 344059) kept on ice. Using a set of glass dounce tissue grinders (Sigma-Aldrich, D8938), a sample of frozen tissue was subjected to dounce homogenization (10 times with loose pestle followed by 10 times with tight pestle) in 1 mL of homogenization buffer (0.32 M sucrose, 5 mM CaCl_2_, 10 mM Tris-HCl pH 8.0, 3 mM MgAC_2_, 0.005% Triton X-100 (Sigma-Aldrich, T9284), 0.1 mM EDTA (ThermoFisher Scientific, AM9260G), 1 tablet of protease inhibitor cocktail). An additional 4 mL of homogenization buffer was added and gently mixed with the homogenate. The mixed homogenate was carefully layered on top of the sucrose cushion in the ultracentrifuge tube. Additional homogenization buffer was added until the total volume of liquid was ~ 13 ml. The tubes were then centrifuged in a Beckman Coulter Optima XPN-100 Ultracentrifuge using a Beckman Coulter SW40 Ti swinging bucket rotor at 25,000 rpm and 4 ℃ for 2 hr. The supernatant was carefully removed via aspiration. The nuclei pellet was incubated on ice for ~ 5 min before it was resuspended in 1.5 mL of chilled resuspension buffer (DPBS (ThermoFisher Scientific, 14190144) and1% BSA (Jackson ImmunoResearch, JAC1000161) and 0.2 U/µL RNase inhibitor (ThermoFisher Scientific, AM2694)). The resuspended nuclei were filtered using a 40-µm cell strainer (BD, 352340) and subsequently transferred to a 1.5-mL tube. The Nuclei were then pelleted at 550 g for 4 min at 4 ℃ and washed with 1.5-mL of resuspension buffer. An additional round of centrifugation and resuspension followed. The Nuclei suspension was visually inspected for morphology and quality assurance and counted using a Nikon Eclipse TE2000 inverted microscope. The suspension was kept on ice before 10× Genomic GEM (gel bead in emulsion) generation.

The volume of nuclei suspension required to generate 10,000 single nuclei GEMs per sample was loaded onto the Chromium Controller (10× Genomics). Libraries were constructed using the Chromium Next GEM Single Cell 3’ Reagent Kits v3.1 (10× Genomics) according to the manufacturer’s specification. Library quantification and quality control were performed using a DNA 1000 chip (Agilengt Technologies), and PE150 sequencing was performed on an Illumina NovaSeq 6000.

Pre-processing with Cell Ranger pipeline

The alignment, filtering, barcode assignment, and UMI counting of paired-end sequencing reads were processed using the “cellranger count” function from the Cell Ranger pipeline (version 5.0.1, 10× Genomics) with the input argument “--transcriptome” set as the file path of mouse reference genome downloaded from the 10× Genomics official website (*https://cf.10xgenomics.com/supp/cell-exp/refdata-gex-mm10-2020-A.tar.gz*).

Downstream processing with Seurat

The CTL-1, CTL-2, STR-1, and STR-2 UMI count matrices were loaded into the Seurat R package (version 4.1.1) [9] to create Seurat objects (genes that expressed in less than 5 cells were filtered out), which were then merged as a single Seurat object. Cells containing less than 1,000 genes or exceeding 5% mitochondrial / 0.05% hemoglobin / 0.05% platelet UMI counts were removed, and only those with novelty scores (the ratio of genes per cell over UMIs) above 0.80 were retained. The filtered Seurat object was split into 4 samples according to the “orig.ident” parameter. The scDblFinder R package (version 1.10.0) [10] was used to identify and exclude potential doublets.

Downstream workflow with SingCellaR

Each sample was individually converted into a “SingleCellExperiment” format via the SingleCellExperiment R package (version 1.18.0), and then loaded into the SingCellaR R package (version 1.2.1)[11, 12] to create a SingCellaR object. Doublet removal was performed using the “DoubletDetection_with_scrublet” function set at default parameters. Cell and gene filtering were performed by assessing QC plots using the “plot_cells_annotation” function. Cells meeting the following QC parameters were included in downstream analyses: 1,000 < UMI counts ≤ 50,000, 500 < number of detected genes ≤ 6,000. Genes expressed in at least 10 cells were included. The effects of UMI counts and percentage of mitochondrial gene expression were regressed out using the “remove_unwanted_confounders” function.

To integrate individual SingCellaR objects, the “SingCellaR_int” object was created. Object file names from individual samples were inputted as the object. The function “preprocess_integration” was performed to combine all UMIs from all samples and cluster together with marker gene information into a single integrated R object. The UMI counts were normalized using the “normaize_UMIs” function with parameter “use.scaled.factor = TRUE”. Highly variable genes were identified using the “get_variable_genes_by_fitting_GLM_model” set at default parameters. Dimensional reduction was performed using the function “runPCA” using unadjusted gene expression values. To perform data integration, the function “runSupervised_Harmony” was run by using 10 principal components (PCs) as determined by the PCA elbow plot generated using the “plot_PCA_Elbowplot” function. After integration, visualization of the data was done using the data embedding method with the function “runFA2_ForceDirectedGraph”, setting the parameters “integrative_method” and “n.dims.use” as “supervised_harmony” and 10 respectively. Clustering analysis was performed using the function “identifyClusters” with “supervised_harmony” as integrative embedding, together with used number of PCs being 10 and local k-nearest neighbor (KNN) equal to 50. Eighteen clusters were identified. To identify marker genes in each cluster, the function “findMarkerGenes” was performed using the “louvain” clustering detection method. According to expression of the marker genes, along with well-established celltype specific gene sets, the 18 clusters were assigned to 11 major celltypes: dorsal neural progenitor cells (NPCd), ventral neural progenitor cells (NPCv), neuroblasts (NB), astrocytes (AC), ependymal cells (EpC), endothilia-like cells (ElC), *Drd1*^+^ spiny neurons (D1), *Drd2*^+^ spiny neurons (D2), oligodendrocyte progenitor cells (OPC), oligodendrocytes (ODC), and microglia (MG).

Cells (except for clusters without neuronal lineage potential, OPC, ODC and MG) expressing at least one of the four cell cycle related genes (*Mki67*, *Top2a*, *Mcm2*, and *Pcna*) were isolated and re-clustered via SingCellaR workflow with above parameters. The potentially dividing cells were clustered into 9 clusters, and annotated as neural progenitor cells (NPC), AC, NB, EpC, ElC, D1, D2, “eccentric” spiny projection neuron (eSPN), respectively.

Cell-cycle position estimation

The cell-cycle position of each potentially dividing cell was estimated using the tricycle R package (version 1.4.0) [13]. Briefly, the “SingCellaR_int” object of the potentially dividing cells was converted into a “SingleCellExperiment” object using the “CreateSeuratObject” function along with the “as.SingleCellExperiment” function of the Seurat R package (version 4.1.1) [9]. The “SingleCellExperiment” object was then projected into the cell cycle embedding of a pre-learned internal reference “neuroRef” via the “project_cycle_space” function of the tricycle R package (version 1.4.0)[13]. The cell cycle position of each cell was assigned via the “estimate_cycle_position” function by the angle formed by PC1 and PC2 in the projected cell cycle space, and then visualized on the force-directed graph embedding using the “plot_emb_circle_scale” function.

Gene regulatory network analysis

The putative gene-to-gene correlated network of the potential dividing cells was inferred using the bigSCale2 algorithm (version 2.0) [14]. Specifically, two normalized gene count matrices generated by the “SCTransform” function from the Seurat package were used to infer the network by default parameters (i.e., clustering = ‘recursive’, quantile.p = 0.9). The network centrality PageRank (PR) was chosen to represent gene essentiality. The network was visualized with the fruchterman-reingold layout using the igraph R package (version 0.7.1). Gene communities in each network were identified using the “cluster_label_prop” function from igraph (version 1.3.4), and clusters consisting of over 5 genes were considered as main communities. Hub genes with maximal biological relevance in each main community were defined as the intersection of three gene sets: 1) top 500 genes with the biggest change in PR absolute value between STR and CTL; 2) genes with over 0.75 quantile of PR value; 3) DEGs between STR and CTL determined by “identifyDifferentialGenes” function of the SingCellaR package (version 1.2.1)[11, 12], set at “min.expFraction = 0.3”. The network’s nodes corresponding to the hub genes of each main community were highlighted in cadetblue (down-regulated in STR) or in darkorange (up-regulated in STR). The hub genes served as the input of GO enrichment analysis using the clusterProfiler R package (version 4.4.4) [7]. Selected hub genes were labeled in the network and listed in the bar plot of the GO enrichment results. The complete list of hub genes can be found in Sup. Table 1.

**Rescue assay.** A MAPK3/1 inhibitor LY3214996 (Selleck, 16 mg/kg) and a MAP2K1/2 inhibitor GSK1120212 (Selleck, 0.35 mg/kg) and the vehicle solution were daily intraperitoneally (IP) injected to 9 to 10-month-old STR male offspring for 27 days (except for the 11^th^ and 22^nd^ days), respectively. Inhibitors were dissolved in the vehicle solution composed of 2% DMSO, 40% PEG300 and 58% ddH_2_O. The injection volume was 200 μl/20g animal. For controls, unstressed male offspring fostered by healthy dams were IP injected with either inhibitors or vehicle solution. Twenty-three hours after the last injection, BrdU was IP administrated, and the animals were sacrificed after 1-hr. A randomization procedure was used to minimize the effects of subjective bias when allocating animals to treatment.

**Lambda Protein Phosphatase (Lambda PP) reaction.** Lambda PP (NEB, P0753S) reaction mixture was prepared using 5 μl of 10× NEBuffer for Protein MetalloPhosphatases (PMP) and 5 μl of 10 mM MnCl_2_, 1 µl of Lambda Protein Phosphatase, supplemented with ddH_2_O to make a total reaction volume of 50 µl. After 3 times of PBS wash (5 min each), cryopreserved sections were immersed in Lambda PP mixture (50 µl/section) for 30 min at 30°C, followed by immunostaining.

**Imaging and Statistical Analysis.** Stained sections were imaged with Zeiss LSM800 confocal microscope as tiled or single images. For sample quantification, we examined sections collected from minimal 3-5 brains of different litters per experimental condition. Only comparable regions between CTL and STR brain sample were imaged and used for quantification. Images obtained were processed and quantified with Imaris software and ImageJ.

Statistical analyses of the data were carried out with R-package, GraphPad Prism, and SAS (Statistical Analysis Software). Mann-Whitney test (non-parametric t-test) was used to assess statistical significance between independent experimental groups. All reported significant levels represent two-tailed values.

**Figure legends**

**Sup.Fig.1.** **Proliferating cells** **in the E-SVZ of lateral wall at different postnatal stages.** CPu, caudate putamen; CP, choroid plexus; LV, lateral ventricle; Sept, septum. Scale bars, 150 μm (left panels), 30 μm (right panels). For sample quantification, we examined sections collected from minimal 3-5 brains of different litters per experimental condition.

**Sup.Fig.2.** **STR female offspring are resilient to PNS.** **(A)** PNS did not significantly increased proliferation of NSC/NPC in the E-SVZ of STR female offspring (P0.5-180). **(B)** Immunostaining of proliferating cells in the E-SVZ. MKI67, green color; PROM1, red color; GFAP, white color. The enlarged images on the right show MKI67/PROM1 double positive ependymal cells are GFAP negative. MKI67^+^/PROM1^+^ cells: CTL = 4.1 ± 0.4%, n = 12; STR = 3.9 ± 0.4%, n=16. P= 0.607. Scale bars, 10μm, 2μm. **(C)** BrdU (green color)/PROM1 (red color)/GFAP (white color) triple immunostaining. The enlarged images on the right show BrdU/PROMl double positive ependymal cells are GFAP negative. BrdU^+^/PROM1^+^ cells: CTL=2.1 ± 0.4%, n=18; STR=2.6 ± 0.5%, n=21. P=0.8070. Scale bars, 10 μm, 2 μm. **(D)** GFAP^+^ cells did not respond to PNS in female offspring (CTL=0.55 ± 0.7%, n=18; STR=0.72 ± 0.6%, n=21. P>0.9999). Mann-Whitney test. All data were presented as mean ± SEM. **p < 0.01.

**Sup.Fig.3. PNS increased the number of proliferating TUBB3^+^** **neuroblasts in male offspring. (A)** TUBB3/MKI67 immunostaining showed that the ratio of neuroblasts increased dramatically in the lateral E-SVZ (MKI67^+^/TUBB3^+^ cells: CTL = 8.6 ± 0.9%, n = 12; STR = 17.7 ± 3.1%, n = 10. P = 0.0032). Scale bars, 10μm. **(B)** GFAP/BrdU double immunostaining in the E-SVZ of lateral wall (CTL = 0.89 ± 0.17%, n = 26; STR = 1.37 ± 0.65%, n = 18. P = 0.6403). Scale bars, 10 μm, 2 μm. Mann-Whitney test. All data were presented as mean ± SEM. **(C, D)** The numbers of neuroblasts in the E-SVZ and in the RMS are not significantly altered in STR female offspring. MKI67^+^/TUBB3^+^ cells: CTL = 12 ± 2.0%, n=5; STR = 13.4 ± 5.0%, n=4. P=0.7302. Scale bar, 10 μm (C). The number of proliferating neuroblasts in the RMS in STR female mice and age-matched CTL. BrdU^+^ cells: CTL =14.4% ± 0.7%, n=34; STR = 13.0% ± 0.7%. P = 0.162. MKI67^+^ cells: CTL =27.1% ± 1.5%, n=34; STR = 26.3% ± 1.5%. P = 0.8841. Scale bar, 100 μm. (D). Mann-Whitney test. All data were presented as mean ± SEM.

**Sup.Fig.4.** **PNS perturbs OB glomerular formation and olfactory behavior tests**. **(A)** Calbindin 2 (CALB2) immunostaining shows disorganized glomerular layer (GL) and slightly increased number of CALB2^+^ cells in the external plexiform layer (EPL) and the granular cell layer (GCL) in the olfactory bulb of STR mice. BrdU labeled cells are in white color. The zoomed-in panels on the right show boxed areas in the left panels. Scale bars: 200 μm (left panels), 15 μm (middle panels), 50 μm (right panels). **(B)** Buried food test showed largely normal olfaction to smell and differentiate nonsocial odors in prenatally stress mice (Latency: CTL, 30.8 ± 5.1s, n=12; STR, 32.2 ± 8.0s, n=14. P=0.6673). **(C)** Prenatally stress mice showed impaired ability to differentiate different social smells (CTL, n=7; STR, n=10). **(D)** Difference of time spent sniffing between social2-1 and social1-3 in (C). Social2-1 - social1-3: CTL, 3.32 ± 0.89s, n=7; STR, -0.39 ± 0.57s, n=10. P= 0.0046. Mann-Whitney test. All data were presented as mean ± SEM. **p < 0.01.

**Sup.Fig.5. Bulk and snRNAseq analyses. (A)** GO analysis shows top GO terms enriched in major gene module (yellow). The top 5 enriched genes of each term are listed. **(B)** Marker gene expression in total sequenced cells and in proliferating cells. **(C)** PNS did not alter overall cell identities in adult offspring. **(D)** Cell cycle position estimation analysis (estimating a continuous cell-cycle pseudotime by principal component analysis of cell-cycle genes).

**Sup.Fig.6. The immunostaining pattern and the specificity of MAPK antibodies. (A)** MAPK3/1 have been shown to interact with many signaling pathways [15-17]. **(B)** To confirm the puncta pattern of p-MAPK3/1 immunostaining, we used a commonly used antibody for unphosphorylated MAKP3/1 (Cell Signaling, 4695). The MAPK3/1 staining showed puncta pattern. Scale bar, 10 μm. The boxed areas were enlarged in lower panels. The cells were highlighted based on nucleus labeling (DAPI). CPu, caudate putamen; LV, lateral ventricle. **(C)** Using a different antibody (Cell Signaling, 4370) against phosphorylated MAKP3/1 to verify the punctate staining pattern. Scale bar, 10 μm. **(D)** Lack of immunoreactivity of p-MAPK3/1 antibody (ThermoFisher, 44-680G) when the brain section was treated with lambda protein phosphatase (lambda PP), suggesting that the staining is specific. Scale bar, 30 μm.
